# Adolescents with major depression featured by sensory-association subtyping show divergent information dynamics and streams

**DOI:** 10.1101/2025.03.29.646114

**Authors:** Xiaobo Liu, Bin Wan, Xihan Zhang, Lang Liu, Siyu Long, Ruiyang Ge, Ruifang Cui, Xin Wen, Guoyuan Yang, Yujun Gao

## Abstract

Adolescent major depressive disorder (MDD) exhibits complex and heterogeneous alterations of brain functional organization. To understand the neurobiological basis of adolescent MDD, we adopted resting-state functional MRI data and used various matrix decomposition approaches to obtain the organization gradients, temporal dynamics, and information streams. With clustering sensory-association gradient features in our exploratory sample (*N*_MDD_ = 250 and *N*_Controls_ = 203), we identified two MDD subtypes. Subtype 1 was characterized by sensory contraction and subtype 2 was associated with association expansion. In addition, two subtypes showed divergent bottom-up and top-down information flows in sensory and association areas using temporal dynamics analysis. These subtypes exhibit distinct age-related changes and reorganization trajectories along sensory-association and auditory-visual axes, highlighting that cortical information flow patterns systematically vary and relate differently to sensory integration, cognitive complexity, and aging. These network distinctions are linked to clinical severity and molecular mechanisms. Subtype 1 is predominantly associated with early neurodevelopmental abnormalities and emotional regulation deficits, while Subtype 2 is more related to synaptic dysfunction and reduced neuronal excitability. These results could be largely replicated in another independent sample (*N*_MDD_ = 73 and *N*_Controls_ = 28). We therefore construct a sensory-association dual functional framework to characterize MDD heterogeneity in adolescent MDD. Itl integrates cortical hierarchy, developmental trajectories, and genetic influences, offering novel insights into MDD pathophysiology and providing a theoretical foundation for precision psychiatry, facilitating personalized diagnosis and intervention strategies.

## Introduction

Adolescent major depressive disorder (MDD) poses significant challenges due to complex manner of clinical symptoms with biological substrates and may exhibit heterogeneous transitions over puberty, resulting in significant challenges in understanding of disease development (Cash et al., 2021; Fox et al., 2013; Sun et al., 2021). These heterogeneous phenomena especially influence the efficacy of pharmacological and psychological interventions, complicating clinical assessments and therapeutic outcomes (Eiland & Romeo, 2013). Neuroimaging, especially functional magnetic resonance (fMRI) provides a noninvasive and objective tool for elucidating the neural substrates underlying this mental variability (Lynch et al., 2024), enabling researchers to identify distinct neurodevelopmental trajectories that may underlie different symptom presentations and treatment responses (Buch & Liston, 2021; Demeter & Greene, 2024; Ecker & Murphy, 2014). Such functional alteration was validated across different samples with various symptom severity and was stable over time, suggesting that intrinsic brain networks serve as heterogeneous neural biomarkers in depression.

Most recently, pioneering work has revealed that large-scale neural networks are systematically organized along multidimensional hierarchical axes, with the sensory-association (S-A) axis as the most representative dimensionality in the intrinsic brain organization (Margulies et al., 2016). This hierarchical axis shifted during adolescence, with progression from lower-order sensory and unimodal regions to higher-order association cortices (Dong et al., 2021; Keller et al., 2023; Luo et al., 2024; Yurgelun-Todd, 2007). Such extended plasticity of late-maturing association cortices is a hallmark of human development (Luo et al., 2024) and a key risk factor for MDD (Xia et al., 2022). For instance, functional gradients in adult MDD patients exhibit significant global and focal alterations, which are associated with sensory processing, higher-order cognitive functions and specific gene expression profiles (M. Xia et al., 2022), indicating adult MDD subtypes could be characterized by their functional deviations (Sun et al., 2023). However, healthy adolescent exhibits significant individual variability in cognitions and their neural representation (Pines et al., 2023a) throughout development (Janssen et al., 2021; S. Zhang et al., 2018), therefore functional heterogeneity of adolescent MDD has not yet been fully elucidated. Here, we aim that adolescent MDD could be classified into distinct neurobiological subtypes along the S-A axis, with different SA axis features having different information interaction dynamics and stream profiles.

Various subtypes of MDD exhibit distinct sensory and association interaction via different temporal dynamic and stream profiles (Frässle et al., 2017, 2021), reflecting how the brain processes external environmental information (Paquola et al., 2025; Park et al., 2024). Adolescent MDDs behaved complex symptoms, encompassing social information processing, cognitive control, and emotion regulation functions and could also be basically summarized by abnormal information processing and exchanging abilities from external environment (Bernaras et al., 2019; Herd & Kim-Spoon, 2021; Ng & Weisz, 2016; Nigg, 2017; Schäfer et al., 2017). Information decomposition theory provides an extending neural framework to elucidate how different MDD subtypes process external information along brain gradients, integrate the diversity of symptoms into a unified cognitive framework, which can even be extended to explain transdiagnostic and comorbid issues in psychiatric disorders (Dalgleish et al., 2020; Fusar-Poli et al., 2019; Jorstad et al., 2023; McQuaid, 2021). Also, the redundant interactions between sensory regions and association regions shape the neurocognitive architecture, offering insights into the neural organization and information processing characteristics of different MDD subtypes (Luppi et al., 2024). For instance, studies have shown that dynamic synergistic measurements derived from fMRI data are significantly disrupted in patients with chronic disorders of consciousness (Luppi et al., 2023), further highlighting the importance of dynamic information processing patterns in psychiatric disorders.

In this study, we leveraged multi-site resting-state functional MRI data to investigate brain reorganization, temporal dynamics, and information stream in adolescents MDD compared to healthy controls. We calculated functional gradients and identified two distinct MDD subtypes using unsupervised machine learning, subsequently analyzing their developmental trajectories. These subtypes were reproducible across independent datasets and exhibited distinct clinical symptom patterns. To further elucidate the aberrant integration patterns between sensory and association cortices, we applied synergy-redundancy information decomposition to analyze the information processing modes associated with functional reorganization in these subtypes.

## Results

### Distinct neurofunctional of two subtypes in adolescent MDD

In this study, we included two independent datasets: an exploratory dataset comprising 258 adolescents MDD (15.18 ± 1.61 years old, 114 females) and 203 healthy controls (15.53 ± 1.98 years old, 82 females), and a validation dataset consisting of 73 adolescents MDD (16.80 ± 1.22 years old, 31 females) and 28 healthy controls (16.82 ± 1.40 years old, 12 females). All participants were aged right-handed, of Han Chinese ethnicity, and free from other psychiatric or major physical illnesses. Each participants’ fMRI data underwent identical quality control procedures, including the exclusion of scans with excessive head motion or image artifacts. Then, each participants’ fMRI time series were then parcellated into Schaefer 400 cortical regions and calculated *Pearson’s* correlation to obtain functional connectivity matrices. Further, the group-level connectivity matrix is obtained by averaging individual functional connectivity matrices. At the group level, connectivity matrices underwent Fisher Z-transformation, retaining only the top 10% of connections per region. A similarity matrix was subsequently generated using the normalized angle kernel, and diffusion mapping was applied to extract functional gradients, yielding a group-level gradient template. The same procedure was applied to individual functional connectivity matrices, and the resulting individual gradients were aligned to the group template via *Procrustes*-based rotation, thereby facilitating consistent comparisons and statistical analyses across groups and subtypes.

Next, we investigated various clustering solutions (from 2 to 10) of adolescent MDD via functional gradient and machine learning algorithms. We observed that the best optimized clustering number was 2 (shown in **Figure 1.a**) and the similarity within subtype 1 (*N =* 135, 15.06 ± 1.54 years old, 58 females/77 males), subtype 2 (*N =* 128, 15.39 ± 2.21 years old, 56 females/67 males) and HC was significantly larger than across subtypes. To further verify the effectiveness of these two subclasses, a support vector machine (SVM) was used to classify the different groups. And we found classification accuracy of subtype 1 and subtype 2 was 93.67%, subtype 1 and HC was 89.98%, subtype 2 and HC was 76.11% (**S-figure 1**). The average functional gradient maps across subtype 1 (*r =* 0.57 ± 0.22), subtype 2 (*r =* 0.65 ± 0.13), and HC (*r =* 0.63 ± 0.18) demonstrated a continuous spatial transition from unimodal to transmodal regions in all groups (**Figure 1.b**). We calculated three global scores for gradients including loading range, individual similarity, and network dispersion (**Figure 1.c**).

**Figure 1.**
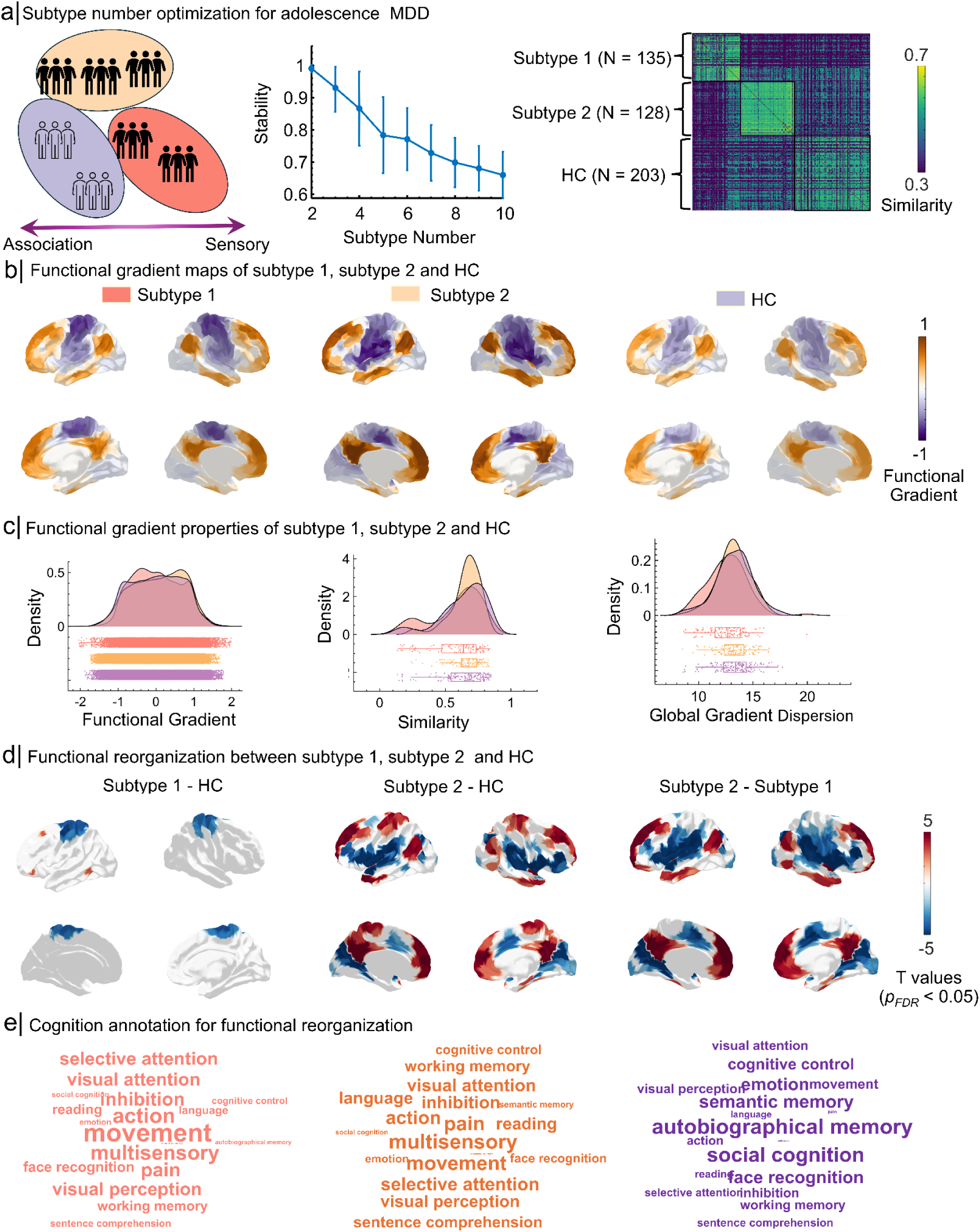
Subtyping adolescent MDD using functional gradients. a) Subtype number optimization for adolescent MDD. The optimal number of subtypes for adolescent MDD was determined to be 2 using stability analysis. Similarity matrices indicate that within-subtype similarity for subtype 1 (*N* = 135), subtype 2 (*N* = 128), and healthy controls (HC; *N* = 158) is significantly higher than across subtypes. b) Functional gradient maps of subtype 1, subtype 2, and HC. Functional gradient maps across subtype 1, subtype 2, and HC show a continuous spatial transition from sensory regions to association regions in all groups. c) Functional gradient properties across groups. Density plots illustrate functional gradient, similarity and global dispersion differences among subtype 1, subtype 2, and HC. Subtype 1 shows lower global gradient dispersion compared to subtype 2 (*t =* -2.71*, p_HSD_ =* 0.003) and HC (*t =* -3.00*, p_HSD_ =* 0.007, with significant distinctions in sensory and association regions. d) Functional reorganization between subtype 1, subtype 2, and HC. Functional reorganization patterns are visualized through group comparisons. Subtype 1 exhibits motor cortex expansion, while subtype 2 shows expansion in associated regions and contraction in auditory and visual regions compared to HC. e) Cognitive annotation for functional reorganization. Cognitive decoding highlights functional differences between subtype 1 and subtype 2. Subtype 1 is associated with sensory functions (e.g., movement and multisensory), while subtype 2 involves higher-order cognitive functions (e.g., reading and inhibition). Word clouds illustrate the cognitive domains linked to functional reorganization patterns in both subtypes.

Specifically, we found subtype 1 shows lower global gradient dispersion compared to Subtype 2 (*t =* -2.71*, p_HSD_ =* 0.0029) and HC (*t =* -3.00*, p_HSD_ =* 0.0074), with significant distinctions in sensory and association regions. We further defined dispersion via calculating *Euclidean* distance between individual regions and manifold centroid to quantify continuous variation along spatial organization through the cortex. Regions exhibiting higher global dispersion values demonstrated greater segregation, whereas regions with lower values showed enhanced integration with proximal areas. Network-level results could be seen in **S-Figure 2**.

**Figure 2.**
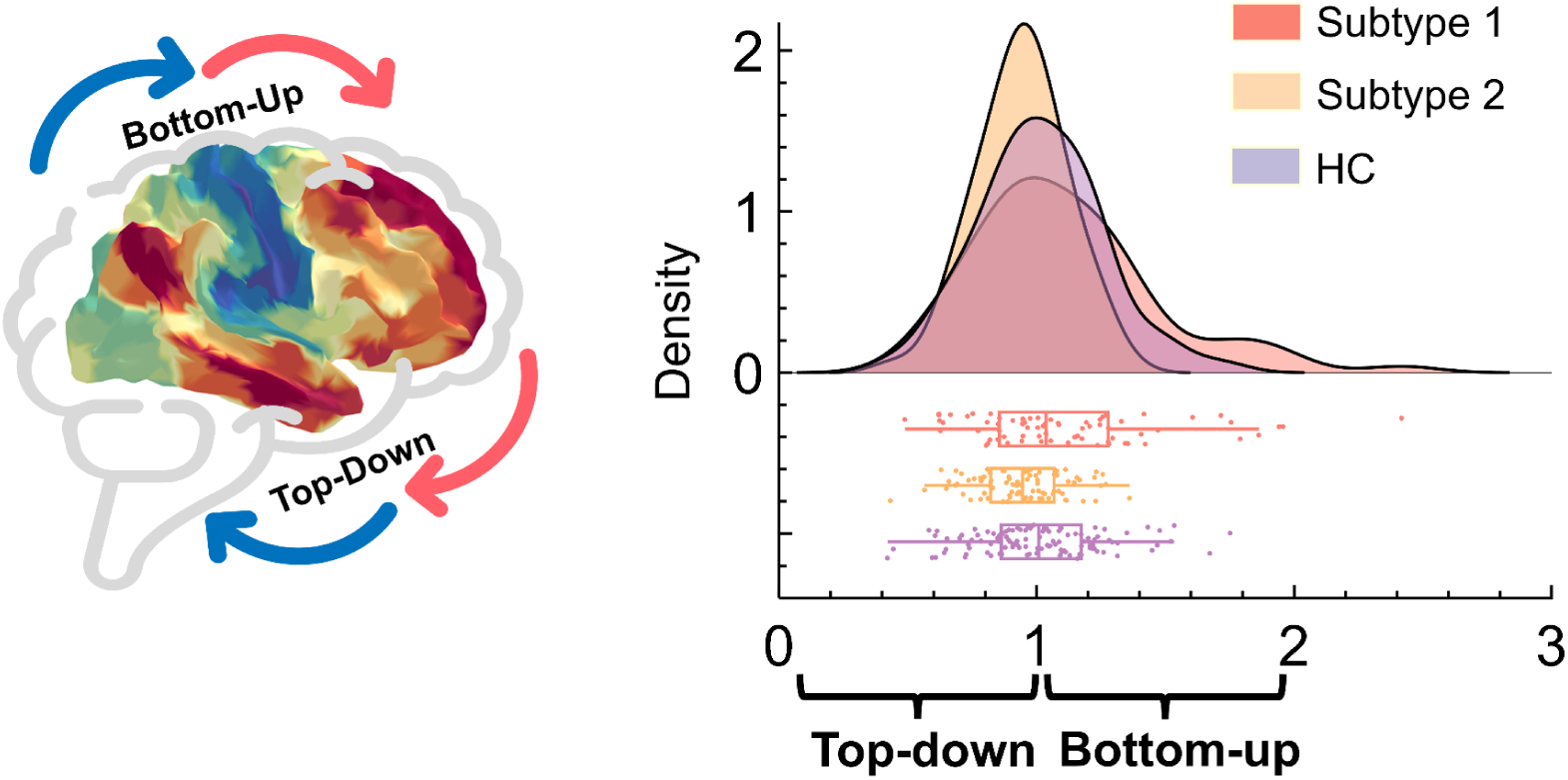
Cortical dynamics flow bottom-up in subtype 1 while top-down in subtype 2. The flowing ratio values in subtype 1 were significantly higher than HC (*t = 3.49, p_HSD_ <* 0.001) and subtype 2 (*t =* 4.68*, p_HSD_ <* 0.001). The flow ratio values in subtype 2 were significantly higher than HC (*t =* - 2.20*, p_HSD_ =* 0.029).

Regarding the regional comparisons (**Figure 1.d**), we found a contraction pattern in the motor cortex in subtype 1, while associated regions showed expansion and auditory and visual areas exhibited contraction in subtype 2, compared to HC (*p_FDR_ <* 0.05). Furthermore, neurosynth-based cognition decoding (shown in **Figure 1.e**) results revealed that the functional reorganization pattern of subtype 1 was sensory functions (e.g., movement and multisensory) while subtype 2 were high order cognitions (e.g., reading and inhibition), which further confirmed our statistics results. These results emphasize one subtype was associated with sensory reorganization another was associated with association reorganization. Together, subtype 1 was associated with sensory reorganization, characterized by motor cortex expansion and increased segregation within the sensorimotor and salience networks, while exhibiting reduced segregation in the limbic and default mode networks. In contrast, subtype 2 was linked to association network reorganization, marked by contraction in auditory and visual regions and enhanced segregation within higher-order networks such as the dorsal attention and control networks, significantly greater than in healthy controls. These findings highlight the pronounced heterogeneity in functional reorganization and neural network architecture across adolescent MDD subtypes.

### Two subtypes reflect different neurodynamics

Next, we investigated the dynamics using the temporal information, as suggested by previous studies (Frässle et al., 2021). Briefly describe the methods (one or two sentences). It captures information flows in various regions in subtype1, subtype 2 and HC (**Figure 2** and **S-figure 3**). Here, we focus on the sensory-association information directions: bottom-up (sensory to association) and top-down (association to sensory). Therefore, we calculated the flowing ratios, defined by the outward degree between sensory regions and association regions. Note that it represents global dynamics flowing bottom up if the ratio value is larger than 1 while top-down if this value is less than 1.

**Figure 3.**
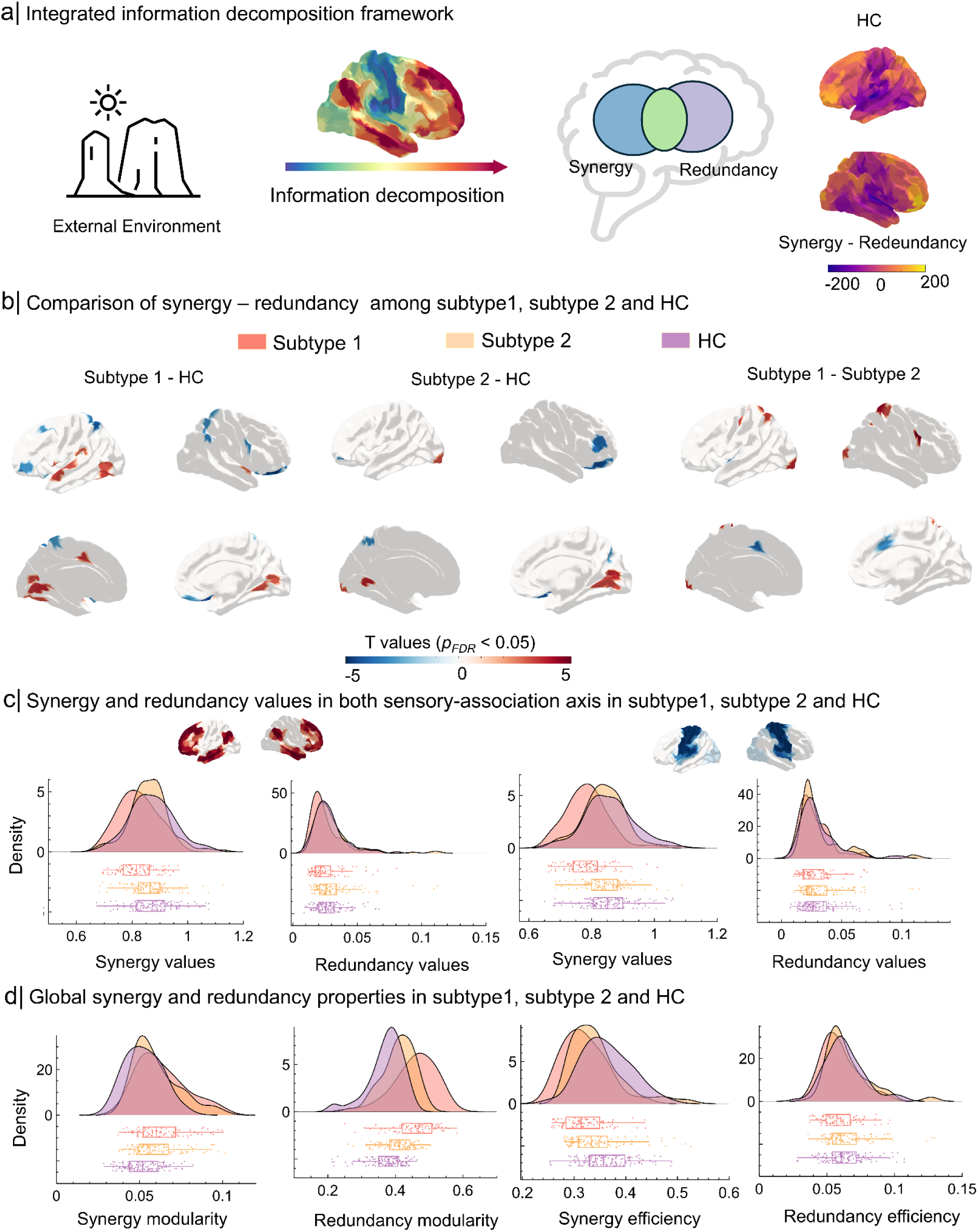
Redundancy-synergy gradients reveal distinct neural information processing patterns in adolescent MDD subtypes. a) Integrated information decomposition framework illustrating synergy (information from regional interactions) and redundancy (shared information across regions) along the sensory-association (S-A) axis. b) Group differences in synergy-redundancy gradients (*p_FDR_* < 0.05). Subtype 1 exhibited elevated sensory synergy compared to subtype 2 and healthy controls (HCs), while subtype 2 showed increased association synergy relative to subtype 1 and HCs. c) Synergy and redundancy values in sensory (left) and association (right) cortices. Subtype 1 demonstrated significantly reduced sensory and association synergy versus Subtype 2 and HCs (*p_HSD_* < 0.001). Subtype 2 showed no synergy differences from HCs. Subtype 1 had lower sensory redundancy than Subtype 2 (*p_HSD_* < 0.001), while subtype 2 displayed higher sensory redundancy than HCs (*p_HSD_* = 0.003). d) Global network properties: Synergy networks in subtype 1 and subtype 2 exhibited higher modularity than HCs (both *p* < 0.001), with subtype 1 showing lower global efficiency than subtype 2 (*p* < 0.001). Redundancy networks in subtype 1 displayed greater modularity and lower efficiency compared to subtype 2 and HCs (*p_HSD_* < 0.001).

We found that the flow ratio values in subtype 1 (*t =* 3.24*, p <* 0.001), subtype 2 (*t =* - 3.68*, p <* 0.001) and HC (*t =* - 0.52*, p >* 0.05) were significantly different from 1 via *one-sample t-*test. Then we further compared the ratio values among subtype 1, subtype 2 and HC via *One-way ANOVA* and *post-hoc analysis* with *Tukey’s Honest Significant Difference* (HSD) with sex and age as covariates. The *post-hoc* analysis showed flowing ratio values in subtype 1 were significantly higher than HC (*t =* 3.49*, p_HSD_ <* 0.001) and subtype 2 (*t =* 4.68*, p_HSD_ <* 0.001). The flowing ratio values in subtype 2 were significantly higher than HC (*t =* - 2.20*, p_HSD_ =* 0.029). Together, those results indicated that the dominant dynamic pattern in subtype 1 was more bottom-up while in subtype 2 was more top-down.

### Various age effects on functional gradient of subtype 1 and subtype 2

The adolescence demonstrates a developmental shift in functional organization, initially characterized by a predominant segregation between sensory and motor systems, which subsequently evolves into a more integrated architecture through interactions with later-maturing regions of the association cortex that underpin higher-order cognitive processes (Y. Xia et al., 2022). Therefore, we further explored how functional gradients in two subtypes altered across ages to further clarify age effects and MDD subtypes (**S-Figure 4**, **S-figure 5** and **S-figure 6**). We found age effects in subtype 1, subtype 2 and HC aligned with functional hierarchy. Specifically, age effects in subtype 1 follow motor/auditory-motor axis (*r =* 0.50*, p <* 0.001), age effects in subtype 2 follow sensory-association axis (*r =* 0.41*, p <* 0.001), and age effects in HC aligned sensory-association axis (*r =* 0.64*, p <* 0.001). This indicates that different MDD subtypes exhibit specificity in age-related changes in functional gradients, showing significant deviations from the normal developmental trajectory.

**Figure 4.**
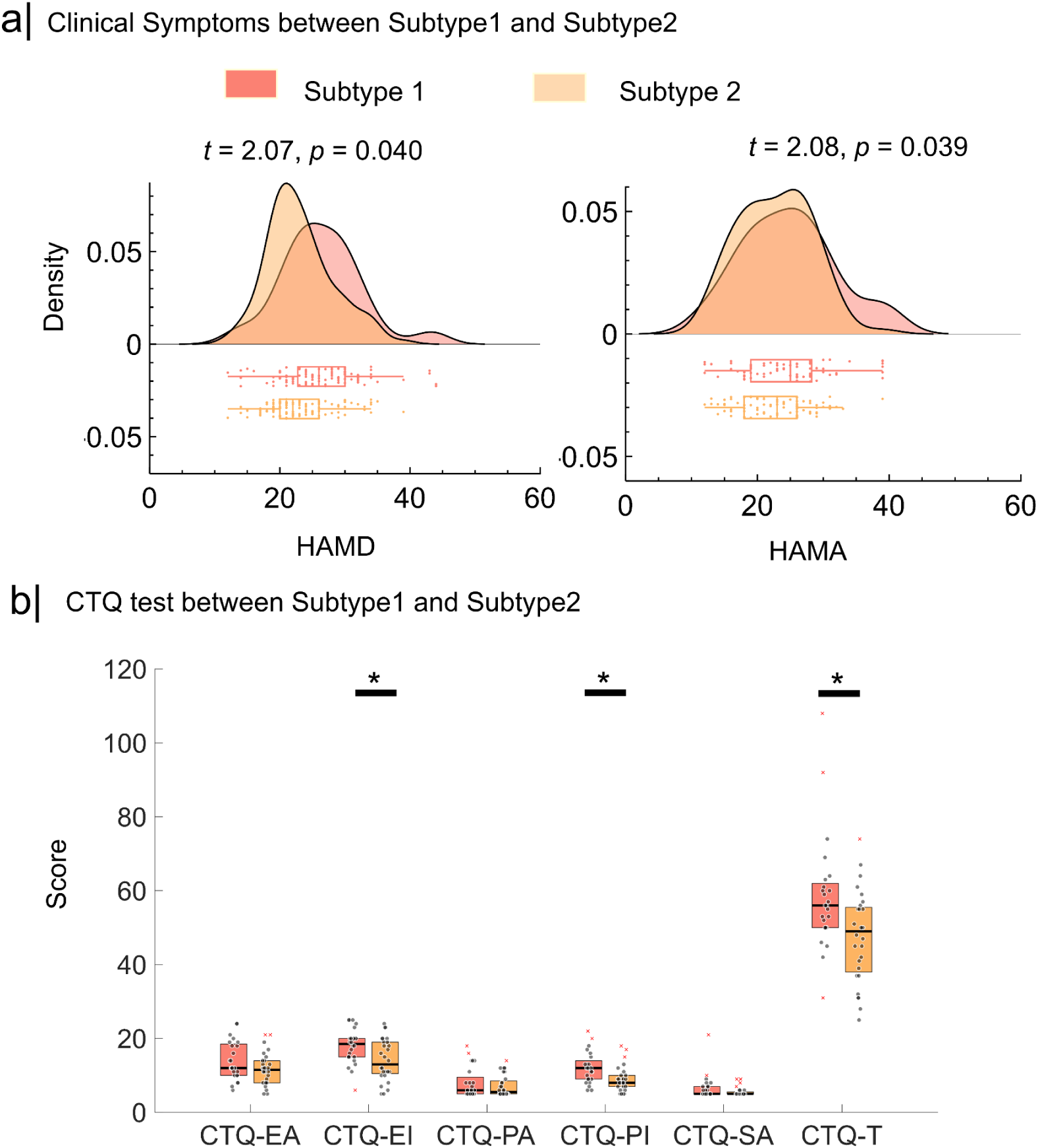
Clinical symptoms and behavior in subtypes 1 and subtype 2. a) Clinical symptoms were significantly different in subtype 1 and subtype 2. Specifically, HAMD (*t =* 2.07, *p =* 0.040) in subtype1 was significantly higher than subtype 2. HAMA (*t =* 2.08, *p =* 0.039) in subtype1 was significantly higher than subtype 2. b) CTQ was significantly different in subtype 1 and subtype 2 (*p_FDR_ <* 0.05). More specifically, CTQ-EI and CTQ-PI in subtype1 were significantly higher than subtype 2 (*p_FDR_ <* 0.05).

**Figure 5.**
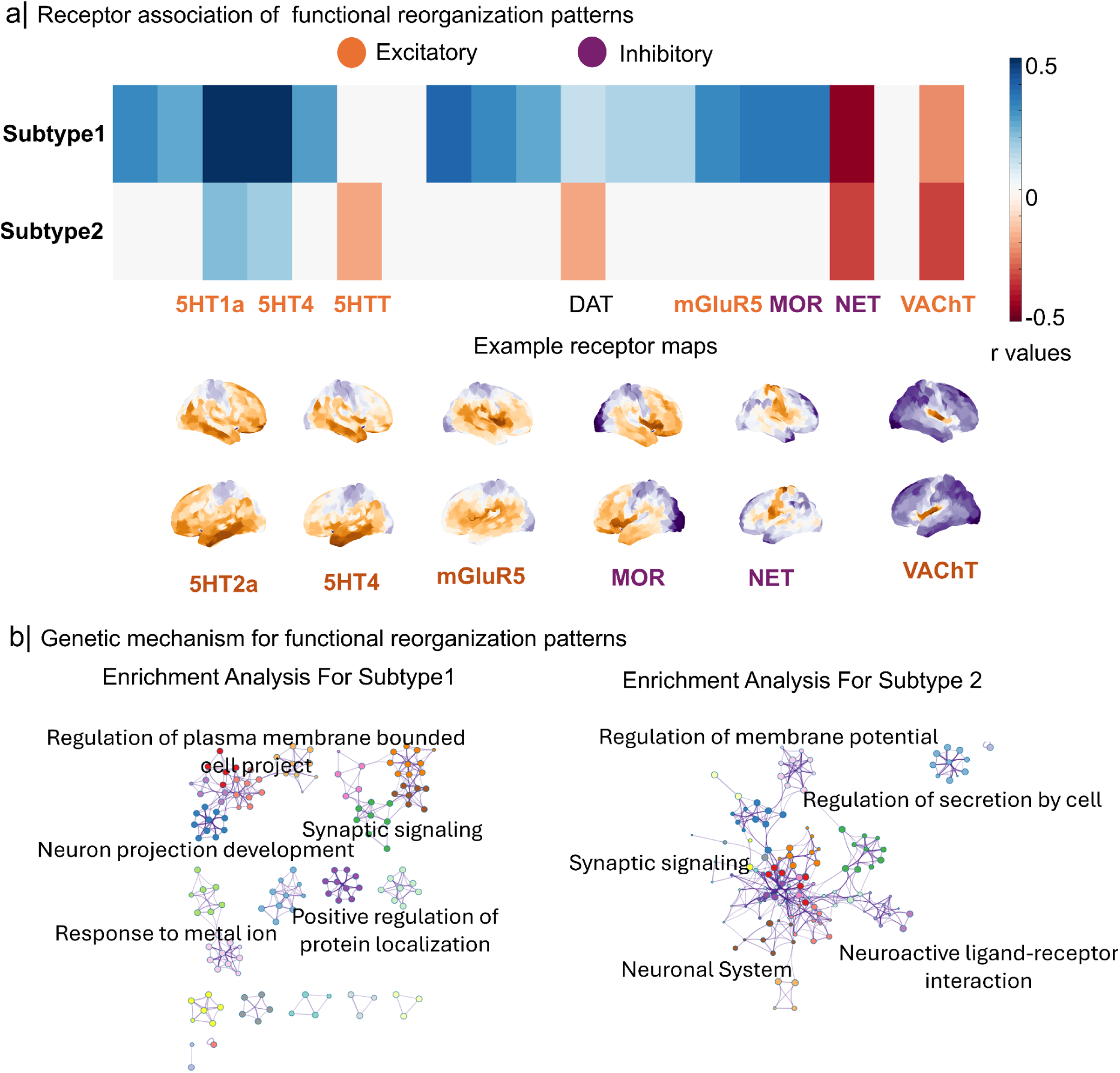
Molecular and Genetic Mechanisms Underlying Functional Reorganization in Adolescent MDD subtypes. a) Spatial correlations between functional reorganization patterns and normative receptor maps revealed distinct associations in subtype 1 and subtype 2. Subtype 1 showed strong correlations with excitatory receptors such as Serotonin Transporter (5HT2a and 5HT4), mGluR5, and Vesicular Acetylcholine Transporter (VAChT), while inhibitory receptors (e.g., GABAa, MOR, and NET) were also implicated. Subtype 2 demonstrated similar receptor associations, emphasizing differential alignment with sensory and association region reorganization. Sensory reorganization correlated with Pax6-expressing cells, Synuclein (Sncg), Astrocytes (Astro), and Layer 5 thick-tufted neurons (L5IT), whereas association area reorganization was linked to Layer 4 thick-tufted neurons (L4IT). b) Gene enrichment analysis highlighted distinct molecular processes for each subtype. Subtype 1 was associated with neuronal growth and structural plasticity, including regulation of plasma membrane-bound cell projections, synaptic signaling, and neuron projection development. Subtype 2 was linked to processes related to synaptic efficiency and neuronal excitability, including regulation of membrane potential, neurotransmitter signaling, and neuroactive ligand-receptor interaction. These findings suggest subtype-specific functional reorganization mechanisms, providing insights into potential molecular targets for therapeutic interventions.

### Redundancy -synergy gradients reveal distinct neural information processing patterns in adolescent MDD Subtypes

Information from the external environment is processed along a gradient toward deeper brain regions (Luppi et al., 2024). Therefore, we applied Integrated Information Decomposition to resting-state fMRI data across different subtypes of MDD (**Figure 3.a**). This methodology allowed us to quantify the extent to which information about the brain’s future trajectory is redundantly encoded by distinct brain regions, as well as the extent to which it is mediated by synergistic interactions between regions. By differentiating between synergistic and redundant interactions in the time series of 400 brain regions, we independently ranked each brain region based on the degree of its synergistic and redundant interactions with other regions. The difference between these rankings (synergy values - redundancy values) was used to determine the relative contribution of each region to synergistic versus redundant processing, thereby establishing a redundancy-to-synergy gradient across brain regions. Statistical analyses show that positive values represent higher synergistic interactions, indicating that brain regions are more inclined to process information through cross-region synergy. In contrast, negative values indicate higher redundant interactions, reflecting a tendency for brain regions to share similar information.

Our results redundancy-synergy gradient (**S-Figure 7** and **S-Figure 8**) aligned with the sensory-association axis in subtype 1 (*r =* 0.50, *p_spin_ <* 0.05), subtype 2 (*r =* 0.40, *p_spin_ <* 0.05) and HC (*r =* 0.55, *p_spin_ <* 0.05). This indicates that the information interactions between brain regions follow the principle of functional hierarchy, consistent with previous findings (Luppi et al., 2024). The figure illustrates the synergy-redundancy distribution across subtype 1, subtype 2, and HC. Subtype 1 exhibited greater synergy in sensory regions, accompanied by lower redundancy in association regions (**Figure 3.b**). Subtype 2 showed significantly higher synergy in association regions compared to HC, along with reduced redundancy in sensory regions. Additionally, subtype 1 displayed higher synergy in sensory regions than subtype 2, while subtype 2 demonstrated greater synergy in association regions compared to subtype 1. The primary feature of subtype 1 is elevated synergistic interactions in sensory regions, highlighting functional abnormalities in sensory processing. In contrast, subtype 2 is characterized by higher synergistic interactions in association regions, potentially linked to disruptions in higher-order cognitive functions.

Furthermore, we compared the sensory and association cortical regions’ synergy and redundancy capabilities across adolescent depression subtypes 1, 2, and HCs (**Figure 3.c and Figure 3.d**). Specifically, we found that subtype 1 exhibited significantly lower synergy in both sensory (*t =* -6.00, *p_HSD_ <* 0.001) and association regions (*t =* -4.25, *p_HSD_ <* 0.001) compared to subtype 2 and healthy controls (sensory regions: *t =* -7.26, *p_HSD_ <* 0.001; association regions: *t =* -5.13, *p_HSD_ <* 0.001). Notably, subtype 2 did not show abnormal synergy compared to healthy controls (sensory regions: *t =* -1.61*, p_HSD_ >* 0.05; association regions: *t =* -0.92*, p_HSD_ >* 0.05). We further analyzed network modularity and global efficiency, where modularity measures the extent of functional segregation between brain regions, and global efficiency reflects the overall capacity for information integration. Subtype 1 exhibited significantly higher modularity than subtype 2 (*t =* 1.56*, p_HSD_ >* 0.05) and HC (*t =* 6.38*, p_HSD_ <* 0.001), while global efficiency was lower than subtype 2 (*t =* -3.70*, p_HSD_ <* 0.001) but not significantly different from healthy controls (*t = -7.18, p_HSD_ > 0.05*). Subtype 2 showed higher modularity (*t =* 4.91*, p_HSD_ <* 0.05) but lower global efficiency (*t =* -3.06*, p_HSD_ =* 0.005) compared to controls. These results suggest that subtype 1 is characterized by greater network segregation and reduced information integration, while subtype 2 maintains better integration capacity but still exhibits reduced efficiency compared to controls.

Similarly, in analyzing redundancy networks, subtype 1 exhibited significantly lower sensory and association redundancy compared to subtype 2 (sensory regions: *t =* -3.39, *p_HSD_ <* 0.001; association regions: *t =* -2.76, *p_HSD_ =* 0.0063), but there were no significant differences compared to controls (sensory regions: *t =* -1.03, *p_HSD_ >* 0.05; association regions: *t =* -0.99, *p_HSD_ >* 0.05). Subtype 2 demonstrated significantly higher sensory redundancy than HC (*t =* 2.96, *p_HSD_ =* 0.003) with no differences in association regions (*t =* 1.91, *p_HSD_ >* 0.05). Further, we found that subtype 1’s network modularity was significantly higher than subtype 2 (*t =* 6.34, *p_HSD_ <* 0.001) and controls (*t =* 11.51, *p_HSD_ <* 0.001), but its global efficiency was significantly lower than subtype 2 (*t =* -4.06, *p_HSD_ <* 0.001) with no significant difference from controls (*t =* -0.95, *p >* 0.05). Compared to controls, subtype 2 had both higher modularity (*t =* 5.97, *p_HSD_ <* 0.001) and global efficiency (*t =* 3.02, *p_HSD_ =* 0.007). These findings indicate that subtype 1 shows weaker integrative capacity in redundancy, whereas subtype 2 exhibits higher sensory redundancy, but its association redundancy is comparable to healthy controls. This suggests that the two subtypes may be underpinned by distinct neurobiological mechanisms and information processing strategies.

In all, subtype 1 exhibits significantly reduced synergy in both sensory and association regions, reflecting increased functional segregation and diminished integrative capacity. Conversely, although subtype 2 demonstrates overall synergistic interactions comparable to healthy controls, it shows significantly elevated redundancy in sensory regions, suggesting possible excessive information sharing among these areas.

### Clinical profiles in two adolescent MDD subtypes

Next, we probed the clinical symptoms in two subtypes (**Figure 4),** including Hamilton Depression Rating Scale (HAMD), Hamilton Anxiety Scale (HAMA) and Childhood Trauma Questionnaire (CTQ). Both HAMD (*N_Subtype 1_ =* 101, *N_Subtype 2_ =* 113, *t =* 2.07, *p =* 0.040) and HAMA (*N_Subtype 1_ =* 73, *N_Subtype 2_ =* 86, *t =* 2.08, *p =* 0.039) of subtype 1 are significantly higher than subtype 2 (**Figure 4.a)**. CTQ was significantly different in subtype 1 and subtype 2 (*N_Subtype 1_ =* 24, *N_Subtype 2_ =* 28, *p <* 0.05). More specifically, CTQ-EI and CTQ-PI (**Figure 4.b)** in subtype1 were significantly higher than subtype 2 (*p_FDR_ <* 0.05).

### Molecular mechanisms and genetic mechanisms of Functional reorganization

Finally, we performed spatial correlation analysis to investigate the alignment between molecular/cellular maps and functional reorganization maps for various adolescent MDD subtypes (**Figure 5.a)**. These normative molecular and cellular maps, derived from the Neuromaps platform (Markello & Misic, 2021; X.-H. Zhang et al., 2023), represent various neurotransmitter and cellular systems. Spatial correlations enabled us to identify which receptor systems and cell types were spatially aligned with the sensory-association reorganization axis across different episodes. Specifically, negative associations indicated that higher cell or neurotransmitter density was more related to sensory reorganization, while positive correlations were associated with the reorganization of association regions.

Similarly, we further showed the functional reorganization of subtype1 and subtype2 was significantly related to excitatory receptors (e.g., Serotonin Transporter (5HT2a and 5HT4), mGluR5 and Vesicular Acetylcholine Transporter (VAChT), inhibitory receptors (e.g., GABAa, MOR NET) and other receptors (e.g., Alpha4Beta4 Nicotinic Acetylcholine Receptor (α4β4)). Figure 6a highlights the molecular and cellular correlations: sensory area reorganization was linked to Pax6-expressing cells, Synuclein (Sncg), Astrocytes (Astro), and Layer 5 thick-tufted neurons (L5IT), whereas association area reorganization correlated with Layer 4 thick-tufted neurons (L4IT). These findings suggest that distinct genes and cell types contribute differentially to the functional reorganization processes in sensory versus association brain regions, potentially reflecting region-specific adaptations.

Also, we implemented enrichment analysis for subtype 1 and subtype 2 (**Figure 5.b)**. Subtype 1 functional reorganization was associated with regulation of plasma membrane bounded cell project, synaptic signaling, neuron projection development, positive regulation of protein localization, and response to metal ion. Subtype 2 functional reorganization was associated with regulation of membrane potential, regulation of secretion by cell, synaptic signaling, neuronal system and neuroactive ligand-receptor interaction. These findings indicate that subtype 1 is primarily associated with broader processes of neuronal growth, connectivity, and synaptic communication. The functional reorganization in this subtype emphasizes structural plasticity (e.g., neuronal projection development) and the cell’s responsiveness to environmental cues (e.g., metal ions). In contrast, subtype 2 focuses on synaptic efficiency and neuronal excitability, particularly related to the regulation of membrane potential and neurotransmitter signaling. This suggests that subtype 2 may exhibit disruptions in synaptic function and communication efficiency, affecting the brain’s ability to transmit signals rapidly and precisely.

Together, these findings reveal that the functional reorganization patterns observed across various BD episodes are closely aligned with the cortical distribution of chemical neuromodulator systems, providing insights into potential molecular biomarkers that could guide future therapeutic strategies.

### Replication of key findings

Finally, we replicated the depression subtypes via an independent CAT-D open dataset (**S-figure.11**). We found the optimization of the subtype number was two. And we also found subtype 1 was associated with motor contraction and subtype 2 was association expansion, which was confirmed by the previous results.

## Discussion

This study identified two distinct subtypes of adolescent MDD: subtype 1, characterized by sensory region reorganization, exhibited a bottom-up information flow pattern, heightened network segregation, and reduced integrative capacity. Subtype 2, associated with association region reorganization, displayed a top-down information flow pattern and better integration, although still below the level observed in healthy controls HC. Both subtypes demonstrated age-related alterations in functional gradients, significantly deviating from normal developmental trajectories. Subtype 1 showed higher synergy in sensory regions and lower redundancy in association regions, whereas subtype 2 exhibited greater synergy in association regions and reduced redundancy in sensory regions. Clinically, subtype 1 presented with more severe symptoms, linked to widespread disruptions in early development and emotional regulation, while subtype 2 was closely associated with impairments in synaptic efficiency and neuronal excitability. In summary, our study proposes a binary hierarchical model systematically characterizing the neurophenotypes of adolescent MDD subtypes, offering critical insights into the unique neurobiological mechanisms driving each subtype. This model captures the complex interplay between sensory and association brain regions, integrates developmental and genetic factors, and underscores its significance for precision psychiatry.

This study identifies two major subtypes of functional reorganization in adolescent MDD. Subtype 1 is characterized by reorganization predominantly within sensory cortices, exhibiting a bottom-up flow of information, higher network segregation, and reduced integrative capacity. Subtype 2, by contrast, is driven by association cortices, showing a top-down flow of information and relatively enhanced integration, albeit still below that of HC. These findings converge with the theoretical framework proposed by Margulies et al. (2016), which highlights a continuous gradient of brain connectivity along the S-A axis and underscores its close links to perception and cognition (Margulies et al., 2016). Furthermore, Luppi et al. (2024) emphasize that sensory regions play a pivotal role in information segregation and robustness, whereas association cortices facilitate integrative and flexible processes (Luppi et al., 2022, 2023, 2024). Our distinction between subtype 1 and subtype 2 strongly supports this conceptual model. Leveraging multimodal data from diverse sites and advanced machine-learning algorithms, we provide refined evidence of the heterogeneous neurobiological underpinnings of adolescent MDD, suggesting that even under a single clinical diagnosis, individuals may exhibit functional reorganization patterns led by sensory or association regions.

The present findings further illuminate distinct developmental paths in the two subtypes. Both subtypes show age-related shifts in functional gradients, yet deviate from the normative trajectories observed in healthy peers. Specifically, subtype 1 follows a gradient from motor/auditory to motor regions, while subtype 2 aligns more closely with the sensory–association axis—diverging from the typical adolescent progression toward higher-order association networks. Prior research has underlined that adolescence is a critical window for the protracted maturation of association cortices (Pines et al., 2023), crucial to advanced cognitive and socioemotional integration (Luo et al., 2024). Our results suggest that subtype 1’s excessive reliance on sensory cortex development might heighten sensitivity to external stimuli, whereas subtype 2 shows a partial “compensatory” enhancement in association areas that nonetheless remains suboptimal. Such discrepancies in the developmental timetable and region-specific maturation processes provide novel insights into the pathophysiology and clinical course of adolescent MDD (Anastasiades et al., 2022; Keshavan et al., 2014).

Our synergy–redundancy analysis indicates that subtype 1 has elevated synergy within sensory regions but reduced redundancy in association areas, whereas subtype 2 exhibits enhanced synergy in higher-order association regions while showing lower redundancy in sensory areas. These data align with the conceptual framework by Luppi et al. (2024), wherein synergy reflects emergent information gained through cross-regional integration, whereas redundancy provides robustness and error tolerance(Luppi et al., 2022, 2023, 2024). Subtype 1’s heightened synergy in sensory cortices may lead to oversensitivity to external stimuli, amplifying affective and stress responses (Canbeyli, 2010, 2022; Panzeri et al., 2001). Conversely, subtype 2’s enhanced synergy in higher-order cognitive circuits suggests an attempt at compensatory regulation of emotional and cognitive functions, although it remains deficient compared to healthy controls (Ibanez et al., 2024; Robison et al., 2020). Such synergy–redundancy differentials support our gradient-based findings and lay a theoretical foundation for precision-targeted interventions.

Clinically, we observed pronounced differences in symptomatology between the two subtypes. Subtype 1 presents more severe depressive and anxious symptom profiles, including higher HAM-D and HAM-A scores, as well as a history of CTQ, particularly in emotional and physical neglect. These findings align with Kaczkurkin et al. (2019) on the susceptibility conferred by early pubertal transitions and heightened psychosocial stress (Kaczkurkin, Park, et al., 2019a, 2019b; Kaczkurkin, Raznahan, et al., 2019), as well as Modabbernia et al. (2021) (Modabbernia et al., 2021), who underscore the enduring impact of adverse early-life events. In Subtype 1, the global disruption in environmental and emotional regulation may foster greater network segregation and diminished integrative capacity(Watt & Panksepp, 2009). Subtype 2, despite showing reductions in synaptic efficiency and neuronal excitability, tends to manifest relatively milder clinical symptoms, potentially reflecting partial compensation in association cortices for inadequate sensory integration (Jernigan et al., 2018; Passarello et al., 2022; Peters et al., 2016; Ruffini et al., 2024). These clinical distinctions underscore the tight link between functional reorganization profiles and disease severity, suggesting that future therapeutic strategies should be tailored to subtype-specific clinical features.

At the molecular and genetic levels, subtype 1’s functional reorganization is linked with gene pathways related to neuronal growth, metabolic regulation, and structural plasticity, while subtype 2 is primarily associated with synaptic transmission, membrane potential regulation, and neurotransmitter interactions. These findings converge with prior evidence from (Wan et al., 2022) and (Hansen et al., 2022a) on the roles of neuromodulatory systems in brain functional organization, and they align with (Luo et al., 2024), which postulates that divergences in gene expression profiles during cortical maturation may underlie susceptibility to psychiatric conditions. The differential molecular mechanisms imply that Subtype 1 may exhibit more pervasive deviations in cellular and synaptic architecture (Sullivan & Geschwind, 2019), whereas subtype 2 shows greater deficits in neuronal processing and signal transmission (Chaudhury et al., 2015; Luscher et al., 2011; Rot et al., 2009). Overall, by integrating functional neuroimaging, developmental, and molecular approaches, this work clarifies the heterogeneous neural mechanisms underlying adolescent MDD and highlights novel directions for precision medicine and personalized interventions—such as the identification of molecular biomarkers to guide pharmacological or other therapeutic strategies. In sum, our findings emphasize the multifaceted nature of adolescent depression across functional and genetic landscapes, offering crucial insight for clinical diagnosis and therapeutic innovation.

## Limitation

Several limitations warrant consideration in the current study. First, we did not account for potential influences of brain structure in our analysis. Second, we did not examine the impact of medication on brain structure and function in individuals with MDD, a common challenge in MDD research that still lacks an ideal solution. Lastly, the receptor, cellular, and gene expression maps used in this study were derived from group-averaged data in previous studies, without accounting for individual variability. Future research should aim to validate these findings by incorporating individual-level differences.

## Conclusion

This study integrates multi-center neuroimaging data and machine learning techniques, providing a comprehensive understanding of the pronounced heterogeneity in adolescent MDD with respect to functional reorganization in S-A regions, deviations from normative developmental trajectories, synergy–redundancy information processing, clinical symptomatology, and gene expression profiles. The findings indicate that different subtypes exhibit distinct patterns of bottom-up or top-down information flow, varying degrees of network segregation and integration, and divergence in molecular pathways, all of which correlate closely with clinical severity and childhood trauma. This body of evidence offers novel perspectives for subtype-based diagnosis and mechanistic inquiries in adolescent MDD, underscoring the pivotal plasticity risks associated with the sensory–association axis and highlighting the importance of incorporating both functional reorganization patterns and molecular signatures in the design of precision treatment strategies. Ultimately, these findings not only deepen our understanding of the etiological heterogeneity and neural underpinnings of adolescent depression but also open new avenues for early clinical detection and targeted interventions.

## Method

### Exploratory Dataset 1

This study received approval from the Ethics Committee of Renmin Hospital, Wuhan University, Wuhan, China, and adhered to the principles of the Declaration of Helsinki (2002). Written informed consent was obtained from all participants and their legal guardians after a thorough explanation of the study’s objectives and procedures.

Participants in the MDD group were experiencing their first depressive episode, with no history of manic or hypomanic episodes, and had not received antidepressant treatment prior to the study. Participants who had a Hamilton Depression Rating Scale (HAMD-24) score greater than 17, were Han Chinese, right-handed, and aged 7–17 years. They had not taken any psychotropic, sedative, or analgesic medications in the month preceding the study. Exclusion criteria included a history of neurological disorders (e.g., epilepsy, multiple sclerosis) or significant physical illnesses (e.g., heart disease, cancer), the presence of other psychiatric disorders or borderline personality disorder, intracranial mass, family history of mental illness or self-harm, severe head trauma resulting in loss of consciousness, substance or alcohol abuse, contraindications to MRI (e.g., pacemakers, metal implants), and an inability to cooperate during MRI scanning. HCs were age- and sex-matched volunteers recruited from the community. Inclusion criteria for the HCs included a HAMD-24 score below 7, Han nationality, right-handedness, and an age range of 12–17 years. Exclusion criteria for HCs mirrored those of the MDD group. More details of data collection and quality control could be seen in *supplementary materials*.

### Replication Dataset 2

We also replicated results of Exploratory Dataset 1 using the Characterization and Treatment of Adolescent Depression (CAT-D) from OpenNeuron (ds004627) (Sadeghi et al., 2022). The details could be seen in *Supplementary materials* as well as (Sadeghi et al., 2022).

After quality control, we included 214 adolescent MDDs and 158 HCs in the exploratory dataset, 55 adolescent MDDs and 35 HCs in the replication dataset in this study (see *Supplementary materials*). The demographics of two datasets were shown in **S-Table 1 and S-Table 2**.

### Data preprocessing, quality control and functional connectome

Raw DICOM files from all datasets were converted to Brain Imaging Data Structure (BIDS) (K. J. Gorgolewski et al., 2016)format using HeuDiConv v0.13.1. Structural and functional preprocessing for the Bipolar dataset were conducted with fMRIPrep 23.0.2 (Esteban et al., 2019) and Numpy (K. Gorgolewski et al., 2011) as its underlying framework. Key anatomical preprocessing steps included intensity normalization, brain extraction, tissue segmentation, surface reconstruction, and spatial normalization. Functional data preprocessing involved head motion correction, slice-time correction, and co-registration. The functional time series were then parcellated according to the Schaefer 400x7 atlas and subjected to a confound removal process via Nilearn, following the “simple” strategy outlined by (Wang et al., 2024). The data quality control could be seen in *Supplementary Data quality control*. This involved high-pass filtering, removal of motion and tissue signals, detrending, and z-scoring. Subsequently, functional connectivity matrices were calculated for each participant using zero-lag Pearson correlation coefficients.

### Functional hierarchy definition

We employed BrainSpace (version 0.1.10; https://github.com/MICA-MNI/BrainSpace) to derive cortex-wide functional connectome gradients, adhering to default parameters. Consistent with prior studies (Margulies et al., 2016), the top 10% of weighted connections per region were retained post z-transformation of the data. An affinity matrix was constructed using a normalized angle kernel to capture connectivity profile similarities between regions. A diffusion map embedding approach (Coifman & Lafon, 2006), a robust non-linear manifold learning technique, was utilized to extract the first functional gradient from the high-dimensional functional connectome for each participant. Individual gradient maps were then aligned to template gradients derived from group average connectome.

### Subtypes definition

To definite subtypes of adolescent MDD, we conduct a data-driven approach on function via machine learning model. Specifically, we applied k-means clustering to analyze the functional gradients in adolescents MDD. The optimal number of clusters was identified based on the clustering stability (Ben-Hur et al., 2001). Specifically, we perturbed the data by sampling 80% of the whole datasets and generated two subsample datasets. We then clustered these two subsample datasets respectively and calculated the stability. Noted that stability was measured by the similarity of two cluster index based overlap samples in these two subsample datasets. Notably, stability was rigorously assessed by averaging stability over performing 1000 iterations on the MDD sample, ensuring the robustness of the identified subtypes. Finally, we repeated the whole process and revised the cluster number to get stability of each cluster number.

To validate the specificity of these network patterns, we employed both similarity matrices across participants and other supervised machine learning techniques to assess their ability to differentiate between depressive subtypes. Specifically, we utilized a Support Vector Machine (SVM) with a radial basis kernel and default parameters to classify subtype1, subtype2, and healthy controls. The data were divided into training and testing sets using leave-one-out cross-validation. Model performance was evaluated using accuracy and the Area Under the Curve (AUC), with the entire process repeated 1000 times to ensure robustness.

To estimate functional reorganization across various MDD subtypes, we implemented One-way ANOVA to compare the functional organization pattern between MDD subtypes and HCs respectively with age and sex as covariates. Then False Discovery Rate (*FDR, q < 5%*) was used to consider the multiple comparisons correction. Furthermore, *post-hoc* analysis with *Tukey’s Honest Significant Difference* (HSD) was implemented to clarify the multiple comparison among various MDD subtypes.

### Dispersion of Functional reorganization

In this study, we used the Schaefer 400-region cortical parcellation as well as its associated Yeo’s seven functional network solution including the visual networks (VN), somatomotor networks (SM), dorsal attention network (DAN), ventral attention network (VAN), limbic (Lib), frontoparietal network (CON), default-mode networks (DMN). We calculated global dispersion as the Euclidean distance between individual regions and the manifold centroid. This metric, which corresponds to eccentricity, provides a quantitative measure of whole-brain integration and segregation. Regions exhibiting higher global dispersion values demonstrated greater segregation, whereas regions with lower values showed enhanced integration with proximal areas. To assess within-network organization, we defined a “Dispersion” metric to assess the degree of gradient dispersion within each functional network. Using the VN, SM, DAN, SAN, LIB, CON, and DMN as boundaries, we averaged the gradient values within each network to establish a central reference. The dispersion was then calculated by measuring the distance between the gradient of each node and this central reference. Finally, we compared the gradient dispersion values across different MDD subtypes and HCs within each resting-state network using *two sample t-tests*.

### Cognition decoding for functional reorganization

We then further estimated the cognition relevance of functional reorganization patterns among various adolescent subtypes using Neurosynth (Yarkoni et al., 2011). We used twenty cognition maps from previous study (Margulies et al., 2016), which represented sensory functions and high order cognition. Then we correlated them with functional gradient difference maps between adolescent MDD subtypes and HC via Pearson’s correlation.

### Top-down and bottom-up information flow in various subtypes

We further explore information flow along the sensory-association axis. Top-down was firstly defined as information flow from association to sensory regions and bottom-up as information flow from sensory regions to association regions (Pines et al., 2023). Information flow was measured by granger causality analysis (Deshpande et al., 2009) and graph theory measurement. Then we calculated the ratio between average outward in sensory regions and association regions respectively. This ratio value represents top-down flow when it is less than 1 while bottom-up when it is larger than 1. Finally, the flow ratio in both MDD subtypes and HCs were compared using one-way ANOVA, with head motion, age, handess and sex as covariates. And then post-hoc analysis with Tukey’s Honest Significant Difference (HSD) was implemented to clarify the multiple comparison among various groups.

### Synergy and redundancy: integrated information decomposition

The Integrated Information Decomposition (ΦID) framework (Luppi et al., 2024) integrates key concepts from Integrated Information Theory (IIT) and Partial Information Decomposition (PID) to parse information flow in neural systems. Specifically, ΦID builds on Shannon’s Mutual Information (MI) by quantifying the dependency between two variables and decomposing the information that two sources provide about a target into unique, redundant, and synergistic components. Synergistic information represents insights derived only when both sources interact, while redundant information reflects insights that are independently contributed by each source (Luppi et al., 2024).This decomposition framework sheds light on how brain regions collectively contribute to future states, highlighting the roles of synergy and redundancy in integrating and coordinating neural processes. To address the limitations of the original IIT measure of integrated information (Φ), which could yield negative values in redundancy-dominated systems, Mediano et al. introduced a revised measure (Φ_R) that incorporates redundancy (Mediano et al., 2021). This revised measure ensures non-negativity and provides a more robust metric for assessing the extent to which “the whole is greater than the sum of its parts,” specifically by focusing on synergistic and transferred information (Mediano et al., 2022). In all, this approach is particularly relevant for understanding brain network dynamics in a more nuanced way, where brain regions are identified as either “gateways,” contributing to integration via synergistic interactions, or “broadcasters,” contributing to the distribution of redundant information.

Therefore, using the ΦID framework on time series data parcellated with the Schaefer 400×7 atlas, we generated for each participant a 400×400 synergy matrix and a 400×400 redundancy matrix, respectively indicating the synergistic and redundant relationships among different brain regions.

### Gradient of redundancy-to-synergy relative importance

In our analysis, we constructed synergy and redundancy networks across regions of interest (ROIs) in the brain and evaluated the relative contribution of each ROI to these interactions. First, we computed the nodal strength for each brain region, which reflects the total sum of its connections in the group-averaged matrix. We then ranked all 400 regions by their nodal strength, assigning higher ranks to regions with stronger connections in both synergy and redundancy networks (Luppi et al., 2024). By subtracting each region’s redundancy rank from its synergy rank, we derived a gradient score that ranges from negative to positive values. A negative score indicates stronger association with redundancy (i.e., a higher redundancy ranking than synergy), while a positive score suggests greater involvement in synergistic processes. This gradient reflects relative differences in the balance of synergy and redundancy within the brain, helping to reveal how individual brain regions participate in either integration or redundancy, depending on the broader network dynamics.

### Synergy and redundancy properties

Modularity and efficiency are critical metrics for understanding the functional roles of synergy and redundancy networks in the brain. Modularity quantifies the degree to which nodes cluster into functional modules, reflecting the organization of network architecture. In synergy networks, high modularity indicates robust inter-regional integration, facilitating global information processing, whereas in redundancy networks, high modularity emphasizes distributed independent contributions, supporting localized functional stability. The reduced modularity may signify disrupted functional segregation, leading to impairments in cognitive and emotional regulation. Efficiency, on the other hand, assesses the brain’s information exchange capacity, with local efficiency capturing short-range information transfer and supporting specific regional functions, while global efficiency measures the capacity for long-range integration across the brain. In MDD, decreased global efficiency suggests impaired integration, whereas increased local efficiency may indicate compensatory mechanisms. Together, modularity and efficiency provide insights into network dysfunction in MDD subtypes, elucidating the balance between integration and segregation and highlighting distinct neural mechanisms (Luppi et al., 2024). Therefore, in this study, we further calculated modularity and efficiency of synergy and redundancy matrices for each individual. Finally, we compared those network properties in both MDD subtypes and HCs using one-way ANOVA, with head motion, age, handess and sex as covariates. And then *post-hoc* analysis with HSD was implemented to clarify the multiple comparison among various groups.

### Age association in various subtypes of adolescent MDD

To further elucidate the age-related effects on functional reorganization patterns across different subtypes of adolescent MDD, we calculated the Pearson correlation between age and the functional gradient in each parcellated brain region for each subtype and the healthy control (HC) group. These age-effect maps were subsequently generated, reflecting how age influences the functional gradient within different brain regions across subtypes. Additionally, we computed the age effects for standard sensory-association and auditory/motor-visual axes, which were defined by the functional connectome from 100 healthy adults in the Human Connectome Project (Elam et al., 2021). These maps provided a comparative framework for examining how adolescent MDD subtypes deviate from normal developmental trajectories, shedding light on the neurodevelopmental dynamics underlying the disorder and its relation to age-specific brain reorganization.

### Clinical symptom association in various subtypes of MDD

To further validate the clinical association of various subtypes, we compared HAMD, Hamilton Anxiety Rating Scale (HAMA) and Childhood Trauma Questionnaire (CTQ) between MDD subtype1 and MDD subtype 2 via *two-sample ttest* respectively, regressing out influence of head motion, sex and handness.

### Enrichment analysis of functional reorganization in subtypes

Then cell type fractions were deconvolved from microarray samples downloaded from Allen Human Brain Atlas (AHBA; http://human.brain-map.org/) (Shen et al., 2012). To study how the functional reorganization is regulated by genes, we combined AHBA and the precise brain connectivity pattern for analysis. Regional microarray expression data were obtained from six post-mortem brains (age = 42.50 ± 13.38 years; male/female = 5/1) with 3702 spatially distinct samples. We used the Abagen toolbox (Markello & Misic, 2021)(https://github.com/netneurolab/abagen) to process and map the data to 400 parcellated brain regions from Schaefer 400 parcellation. We further correlated the two subtypes’ functional reorganization maps with those gene expression cortical patterns.

Then we tested whether transcriptionally dysregulated genes in postmortem brain tissue of messenger RNA are expressed most in cortical regions that are associated with gradient abnormalities across different subtype1 and subtype 2. Metascape analysis (https://metascape.org/gp/index.html#/main/step1) automated meta-analysis tools to identify and interpret common or unique pathways across 40 independent knowledge bases (Zhou et al., 2019). The most significantly functional reorganization patterns related genes were input into the Metascape website, and the obtained enrichment pathways were thresholded for significance at 5%, corrected by the FDR and null modal test.

### Receptor mechanism of functional reorganization of various subtypes

Receptor densities were assessed through the utilization of PET tracer investigations encompassing a total of 18 receptors and transporters spanning nine neurotransmitter systems. This data, recently shared by Hansen and colleagues (https://github.com/netneurolab/hansen_receptors) (Hansen et al., 2022a). The neurotransmitter systems include dopamine (D1, D2 DAT), norepinephrine (NET), serotonin (5-HT1A, 5-HT1B, 5-HT2, 5-HT4, 5-HT6, 5-HTT), acetylcholine α4β_2_, M1, VAChT), glutamate (mGluR5), GABA (GABAa), histamine (H3), cannabinoid (CB1), and opioid (MOR)(Markello & Misic, 2021). Volumetric PET images were aligned with the MNI-ICBM 152 nonlinear 2009 (version c, asymmetric) template. These images, averaged across participants within each study, were subsequently parcellated into the Schaefer 200 template. Receptors/transporters exhibiting more than one mean image of the same tracer (5-HT1B, D2, VAChT) were amalgamated using a weighted average (Hansen et al., 2022a).

### Null model

In our current investigation, we sought to delineate the topographic correlations between common-specific network patterns and other salient neural features. To derive robust inferences, we implemented a null model designed to systematically disrupt the relationship between two topographic maps while maintaining their inherent spatial autocorrelation (Hansen et al., 2022b). Initially, we shuffled receptor maps and assessed their relationships with common-specific network patterns. These spatial coordinates formed the foundation for generating null models, achieved by performing randomly sampled rotations and reassigning node values to the nearest resulting parcel. This procedure was repeated 1000 times to ensure robustness. Critically, the rotational transformation was first applied to one hemisphere and subsequently mirrored onto the contralateral hemisphere. The *95th* percentile of occurrence frequencies from both spatial and temporal null models was designated as the threshold for statistical significance. This rigorous thresholding approach allowed us to identify meaningful topographic correlations while controlling for the intrinsic spatial autocorrelation present in the data.

## Data Availability

The clinical data could be accessed according to reasonable requests for corresponding authors. Allen Human Brain Atlas (AHBA; http://human.brain-map.org/). Neuromap (https://netneurolab.github.io/neuromaps/usage.html), ENIGMA toolbox (https://enigma-toolbox.readthedocs.io/en/latest/pages.html) could be download online. Characterization and Treatment of Adolescent Depression (CAT-D) from OpenNeuron (https://openneuro.org/datasets/).

## Acknowledgments

Xiaobo Liu is supported by the China Scholarship Council. Bin Wan is supported by International Max Planck Research School on Neuroscience of Communication: Function, Structure, and Plasticity (IMPRS NeuroCom), Graduate Academy Leipzig, and Mitacs Globalink Research Award. This work was supported in part by the Health of Hubei Province Scientific Research Project under Grant 2020Cfb512, and by the Mental Health Research Institute of Three Gorges University: YCXL-23-11.

## Exploratory Dataset 1

This study was conducted with the approval of the Ethics Committee of Renmin Hospital, Wuhan University, Wuhan, China, adhering strictly to the principles of the Declaration of Helsinki. Written informed consent was obtained from all participants and their legal guardians after they received a comprehensive explanation of the study’s purpose and procedures.

Participants were recruited through the Center of Prevention and Management of Depression at Renmin Hospital, using a combination of advertisements and referrals. Data collection occurred from May 4, 2018, to December 30, 2023. The sample comprised 314 adolescents diagnosed with major depressive disorder (MDD) according to DSM-IV criteria, confirmed by two board-certified psychiatrists using the Structured Clinical Interview for DSM-IV Axis I Disorders (SCID). Participants were aged 14 to 18 years, had an MDD duration of less than 12 months, and had no history of depression-related medication or electroconvulsive therapy. Exclusion criteria included comorbid psychiatric disorders, history of head trauma with loss of consciousness, somatic illnesses affecting brain morphology, substance abuse, left-handedness, serious physical or neurological conditions, contraindications for MRI, and recent medication use within the last five half-lives. MDD severity was assessed with the 17-item Hamilton Rating Scale for Depression (HRSD), and a score of ≥17 was required for inclusion.

253 healthy controls (HCs) matched for age, sex, ethnicity, education level, and handedness were randomly recruited from the local community, with healthy status confirmed using the SCID-I/NP.

Resting-state functional MRI (rs-fMRI) scans were performed on a 3.0T GE scanner at the PET Center of Renmin Hospital, following established protocols. Participants were instructed to lie supine, eyes closed, and remain awake and motionless. Echo Planar imaging (EPI) was employed with TR/TE of 2000/30 ms, 32 slices, 64x64 matrix, 90° flip angle, 24 cm field of view, 3.0 mm slice thickness with no gap, and 212 scans over 16 minutes.

## Magnetic Resonance Imaging Acquisition

Resting-state functional MRI (rs-fMRI) scans were obtained using a 3.0T General Electric scanner at the PET Center of Renmin Hospital of Wuhan University, following previously established protocols. Participants were instructed to lie in the supine position with their eyes closed, remaining awake and as motionless as possible. An echo planar imaging (EPI) sequence was employed with the following parameters: repetition time/echo time (TR/TE) of 2000/30 ms, 32 slices, 64x64 matrix, 90° flip angle, 24 cm field of view, 3.0 mm slice thickness with no gap, and axial scanning performed 212 times over 16 minutes.

## Replication Dataset 2

We also replicated results of Exploratory Dataset 1 using the Characterization and Treatment of Adolescent Depression (CAT-D) from OpenNeuron (Markiewicz et al., 2021) (dataset id is ds004627).

The CAT-D study is an ongoing longitudinal investigation recruiting adolescents through referrals from community practitioners, self-referrals, and advertisements primarily within Maryland, Virginia, and the District of Columbia. Eligible participants must be between 11 and 17 years of age at the time of enrollment, with a history or current diagnosis of major depressive disorder (MDD) or subthreshold depression (s-MDD), or they must be healthy volunteers (HVs). As of March 11, 2021, a total of 279 participants had been enrolled in the CAT-D study. To facilitate specific analyses, additional inclusion criteria were applied to generate subsamples. In this study, we focused on adolescents identified as HVs or those with a history or current diagnosis of MDD, based on clinician ratings using the Kiddie Schedule for Affective Disorders and Schizophrenia–Present and Lifetime Version (KSADS-PL). Moreover, to compare measures pre- and post-COVID-19 pandemic, we included only participants with data available both before and during the pandemic. The World Health Organization’s declaration of COVID-19 as a global pandemic on March 11, 2020, was used as the reference point for our analyses. Pre-pandemic assessments were defined as those conducted between March 11, 2019, and March 10, 2020, while pandemic assessments were defined as those conducted between March 11, 2020, and March 11, 2021. A total of 166 participants were included to address the study’s first objective regarding the association between the pandemic and symptoms of depression and anxiety. These participants had at least one assessment of depression and/or anxiety symptoms before the pandemic and at least one assessment during the pandemic.

## Data quality control

We excluded participants with excessive head motion (threshold >2mm) or without demographic or imaging data. For the exploration dataset, 44 adolescent MDD and 45 HCs were excluded from the data sample. For the replication dataset,16 participants were excluded resulting in a final sample of 101 subjects.

## Partial information decomposition

Shannon’s mutual information (MI) (Luppi et al., 2022, 2023, 2024) quantifies the degree of interdependence between two random variables, X and Y. This relationship is expressed as:

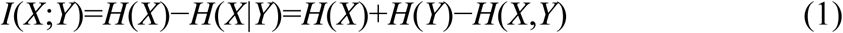

where H(X)H(X)H(X) represents the Shannon entropy of X. The first equality shows that MI measures the reduction in uncertainty (entropy) about X when Y is known, capturing how much information one variable provides about another.

Building upon this concept, (Williams & Beer, 2010) recognized that the information two source variables, XXX and YYY, convey about a third variable, ZZZ, can be further decomposed into unique, redundant, and synergistic components. This led to the development of the Partial Information Decomposition (PID) framework, as formalized in the equation:

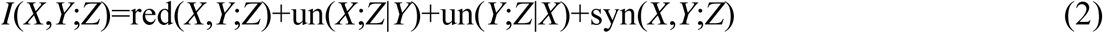

Here, “red” denotes redundancy, or the shared information from both sources, “un” refers to unique information provided by one source but not the other, and “syn” represents synergy, information that can only be extracted when X and Y are considered together.

A classical example of synergy can be demonstrated using the XOR function, where Z=XOR(X,Y)Z = XOR(X, Y)Z=XOR(X,Y). In this system, both X and Y are independent, meaning that neither provides information about Z individually. However, when considered jointly, they fully determine Z, illustrating pure synergy.

While PID offers a robust framework for decomposing information, it leaves open how these components should be measured. For continuous Gaussian variables, most decomposition approaches converge on a solution known as Minimum Mutual Information PID (MMI-PID). This approach defines redundancy based on the minimum mutual information between individual sources and the target, while synergy is quantified as the extra information contributed by the weaker source after considering the stronger source.

Linear Gaussian models are commonly used for functional MRI data as they provide a suitable representation of such time series without the need for more complex nonlinear models. In this study, we adopt the MMI-PID approach, which has been applied in prior neuroscientific analyses. Nonetheless, we also confirm the robustness of our findings by verifying them with discrete data, ensuring that the results do not rely solely on the assumption of Gaussianity.

Though information decomposition is an evolving field, the current approach offers a valuable framework for parsing information flow in neural systems. Further advancements are likely to refine how we compute and interpret these measures, providing deeper insights into brain function.

Finally, segregation and integration coefficients were calculated for each individual and compared between various MDD subtypes and controls using one-way ANOVA, with age, headmotion and sex as covariates.

**S-figure 1.**
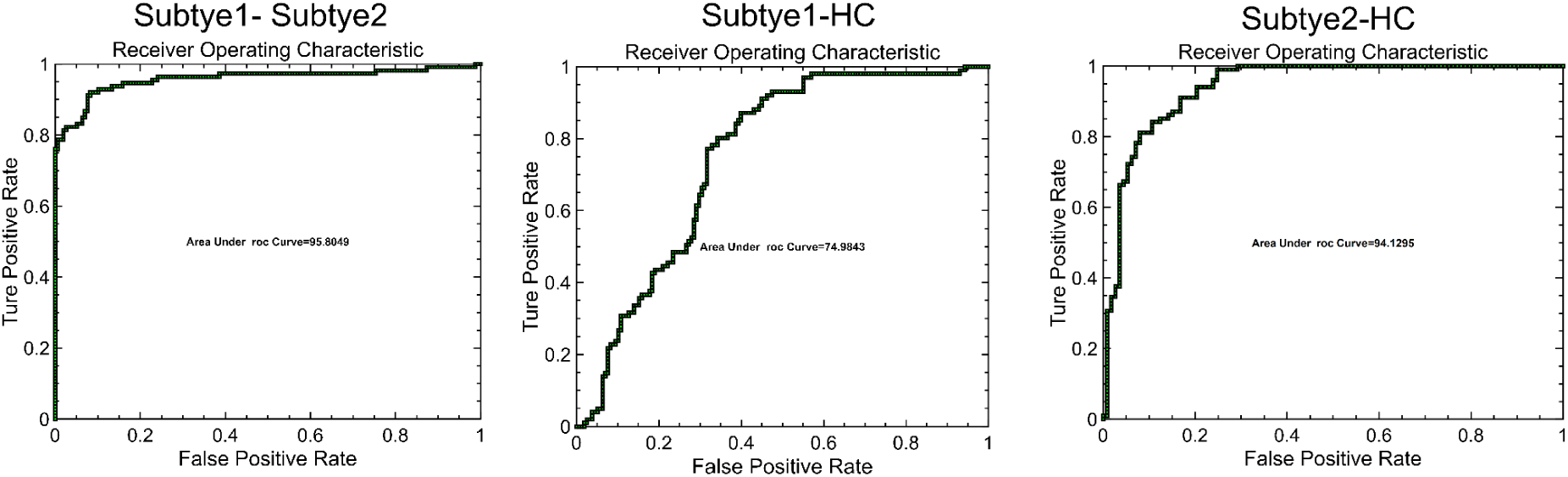
Classification ROC among subtype1, subtype2 and HC. Specifically, the accuracy between subtype 1 and subtype 2 was 93.67% (AUC was 95.81%), subtype 1 and HC was 89.98% (AUC was 74.99%), subtype 2 and HC was 76.11% (AUC was 94.13 %).

**S-figure 2.**
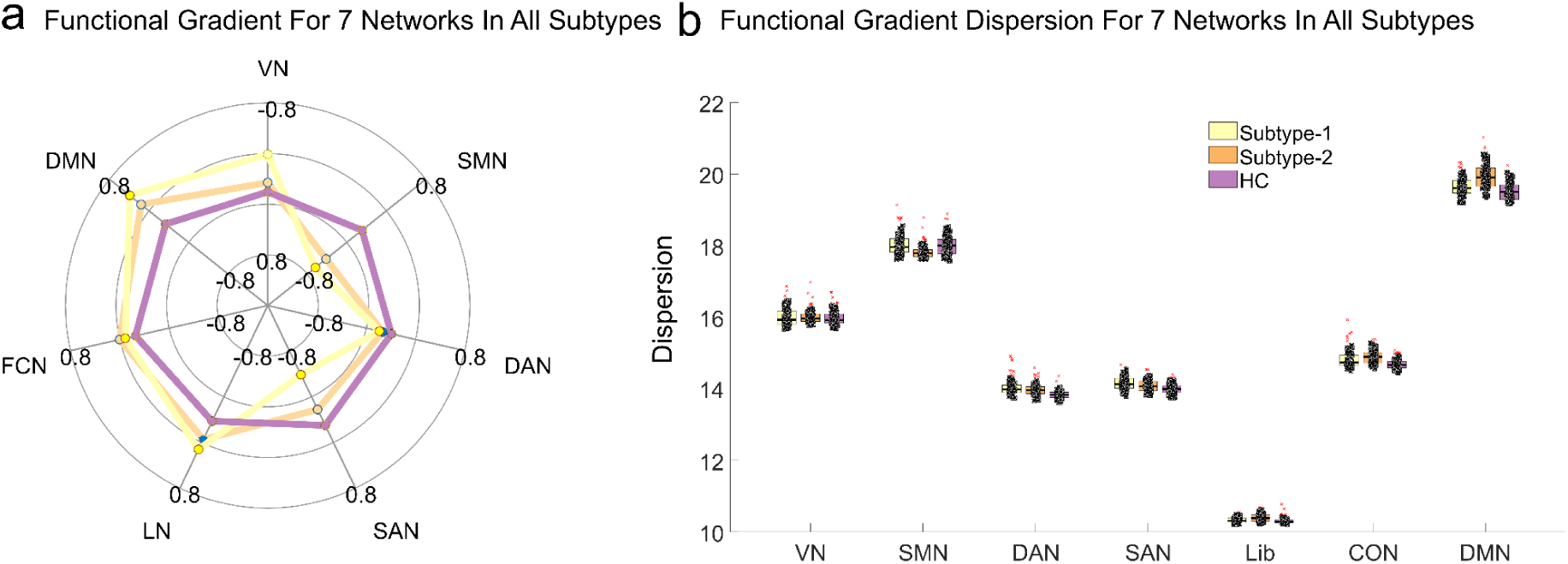
Dispersion of functional reorganization in two subtypes. The results revealed that global dispersion of subtype 1 had significantly lower global dispersion than subtype 2 (*t = -2.75, p_HSD_ < 0.05*) and HC (*t =* -3.10*, p_HSD_ < 0.05*). There is not any significant difference between subtype 2 and HC (*t = -0.29, p_HSD_ > 0.05*). We observed that subtype 1 exhibited significantly higher dispersion in the sensorimotor network (SMN) (*t = 4.82, p_HSD_ < 0.001*) and salience network (SAN) (*t = 2.21, p_HSD_ = 0.028*), but lower dispersion in the limbic network (LIB) (*t = -4.37, p_HSD_ < 0.001*) and default mode network (DMN) (*t = -5.88, p_HSD_ < 0.001*) compared to Subtype 2. Furthermore, when comparing Subtype 1 to healthy controls, we found higher dispersion in the dorsal attention network (DAN) (*t = 7.82, p_HSD_ < 0.001*), salience network (SAN) (*t = 6.24, p_HSD_ < 0.001*), control network (CON) (t = 4.74, *p_HSD_*< 0.001), and default mode network (DMN) (*t = 3.40, p_HSD_ < 0.001*). Similarly, Subtype 2 showed lower dispersion in the sensorimotor network (SMN) (*t = -4.29, p_HSD_ < 0.001*) but higher dispersion in the dorsal attention network (DAN) (*t = 7.45, p_HSD_ < 0.001*), salience network (SAN) (*t = 4.06, p_HSD_ < 0.001*), limbic network (LIB) (*t = 5.92, p_HSD_ < 0.001*), control network (CON) (*t = 8.48, p_HSD_ < 0.001*), and default mode network (DMN) (*t = 10.05, p_HSD_ < 0.001*) compared to healthy controls. No significant differences were observed between the three groups in the visual network (VN) (*p_HSD_ > 0.05*).

**S-figure 3.**
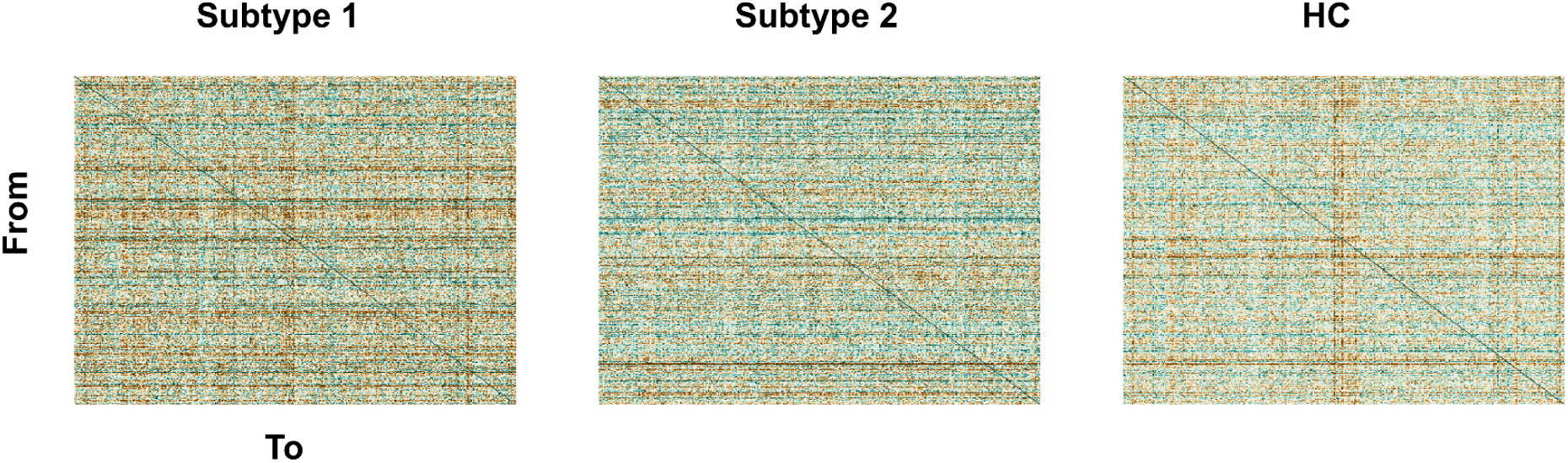
Average GCA matrix for subtype1, subtype 2 and HC.

**S-figure 4.**
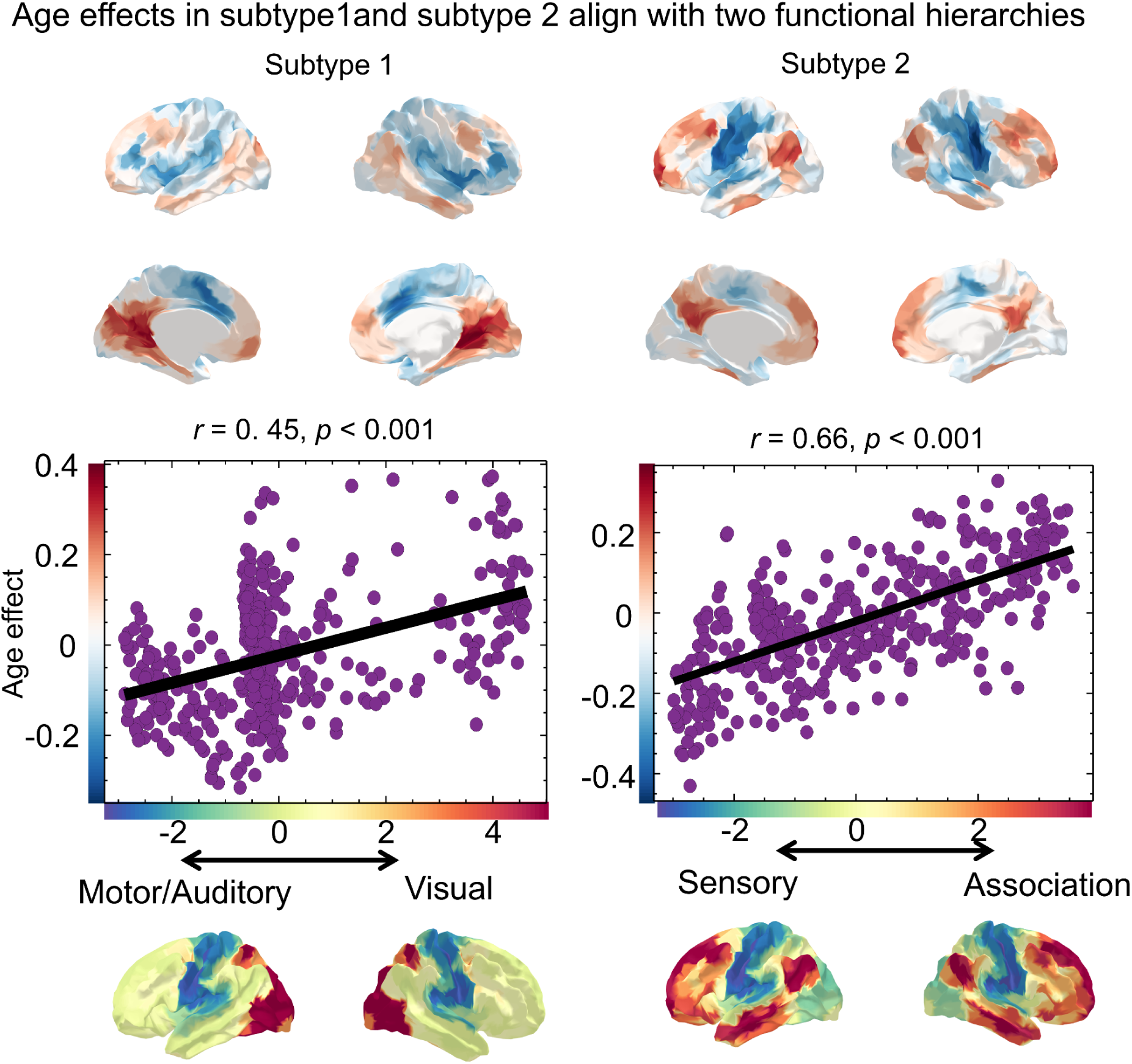
Age effects in subtype 1, subtype 2 and HC aligned with two functional hierarchical axes. Specifically, age effects in subtype 1 follow motor/auditory-motor axis (*r =* 0.50*, p_spin_ <* 0.05) and in subtype 2 follow sensory-association axis (*r =* 0.41*, p_spin_ <* 0.05).

**S-figure 5.**
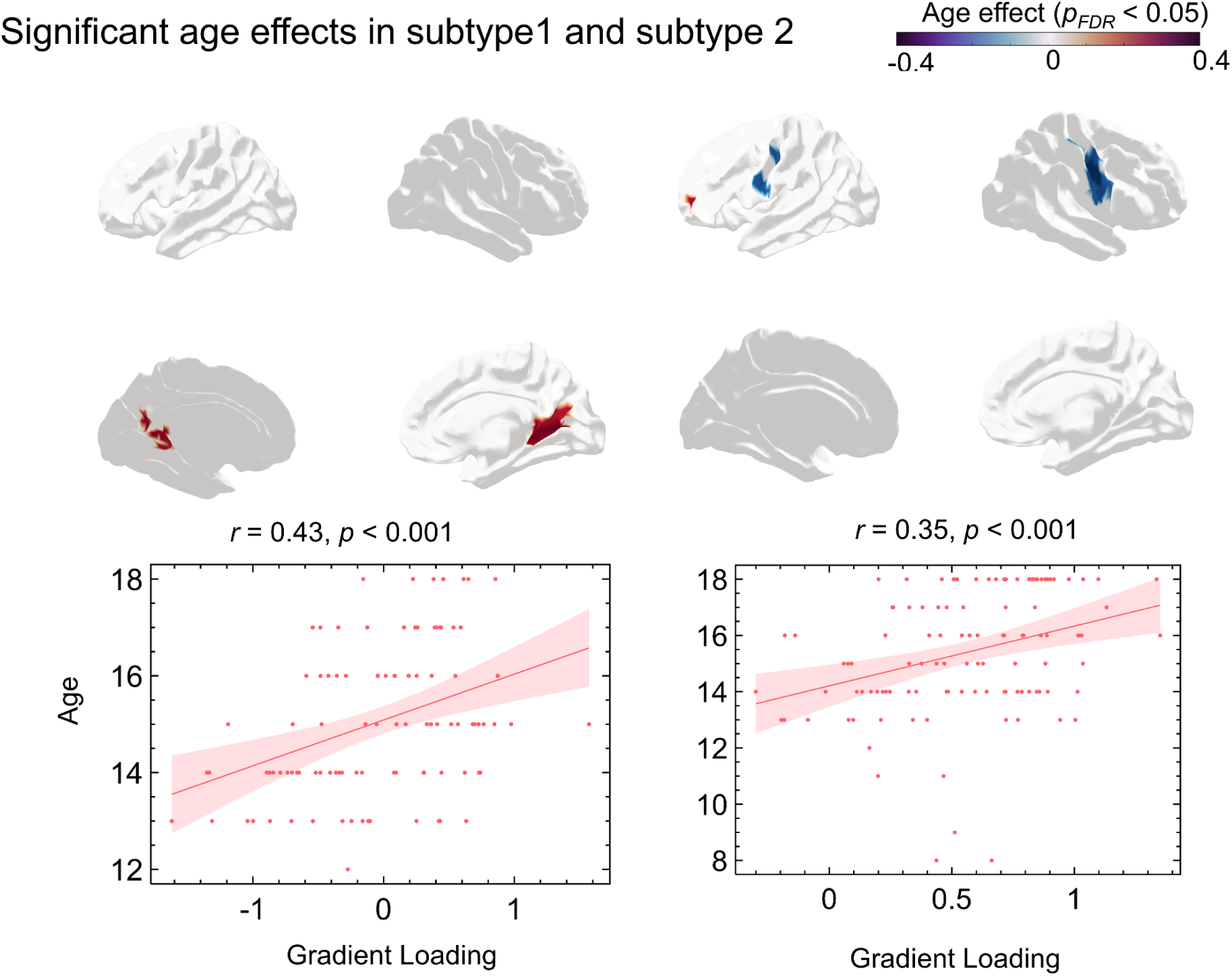
Significant age effects in subtype 1 (*r_peak_ = 0.43, p < 0.001*) and subtype 2 (*r_peak_ = 0.35, p < 0.001*).

**S-figure 6.**
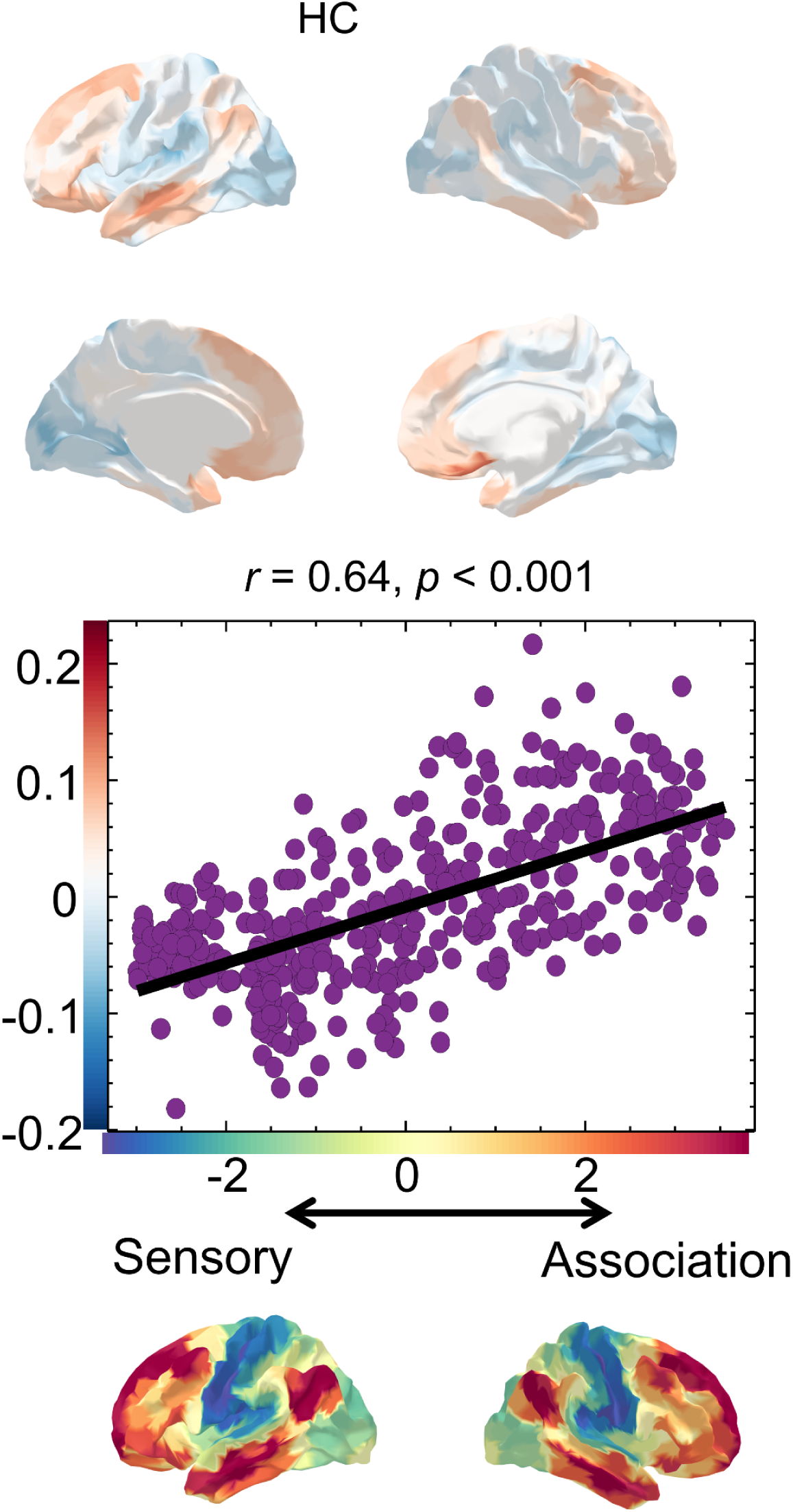
Age effects in HC aligned with the sensory-association axis (*r = 0.64, p < 0.001*).

**S-figure 7.**
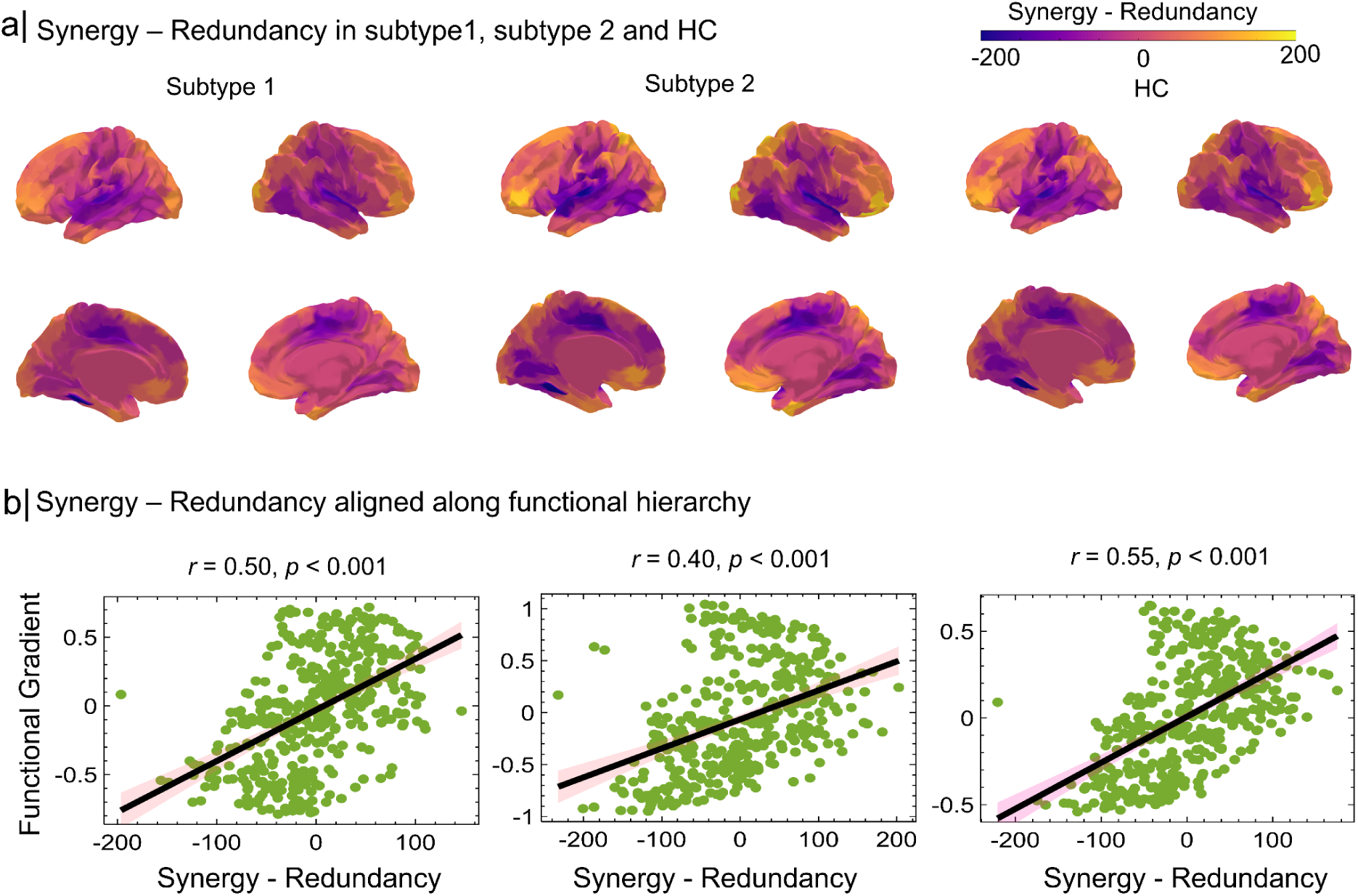
Synergy-Redundancy Gradient in Subtype 1, Subtype 2, and HC. a) The synergy-redundancy gradient maps demonstrate the spatial distribution of synergy (positive values) and redundancy (negative values) across Subtype 1, Subtype 2, and HCs. Subtype 1 exhibits elevated synergy in sensory regions with reduced redundancy in association regions, whereas Subtype 2 shows higher synergy in association regions and lower redundancy in sensory regions compared to HC. b) Synergy-Redundancy aligned with the sensory-association axis in subtype 1 (*r = 0.50, p_spin_ < 0.05*), subtype 2 (*r = 0.40, p_spin_ < 0.05*) and HC (*r = 0.55, p_spin_ < 0.05*).

**S-figure 8.**
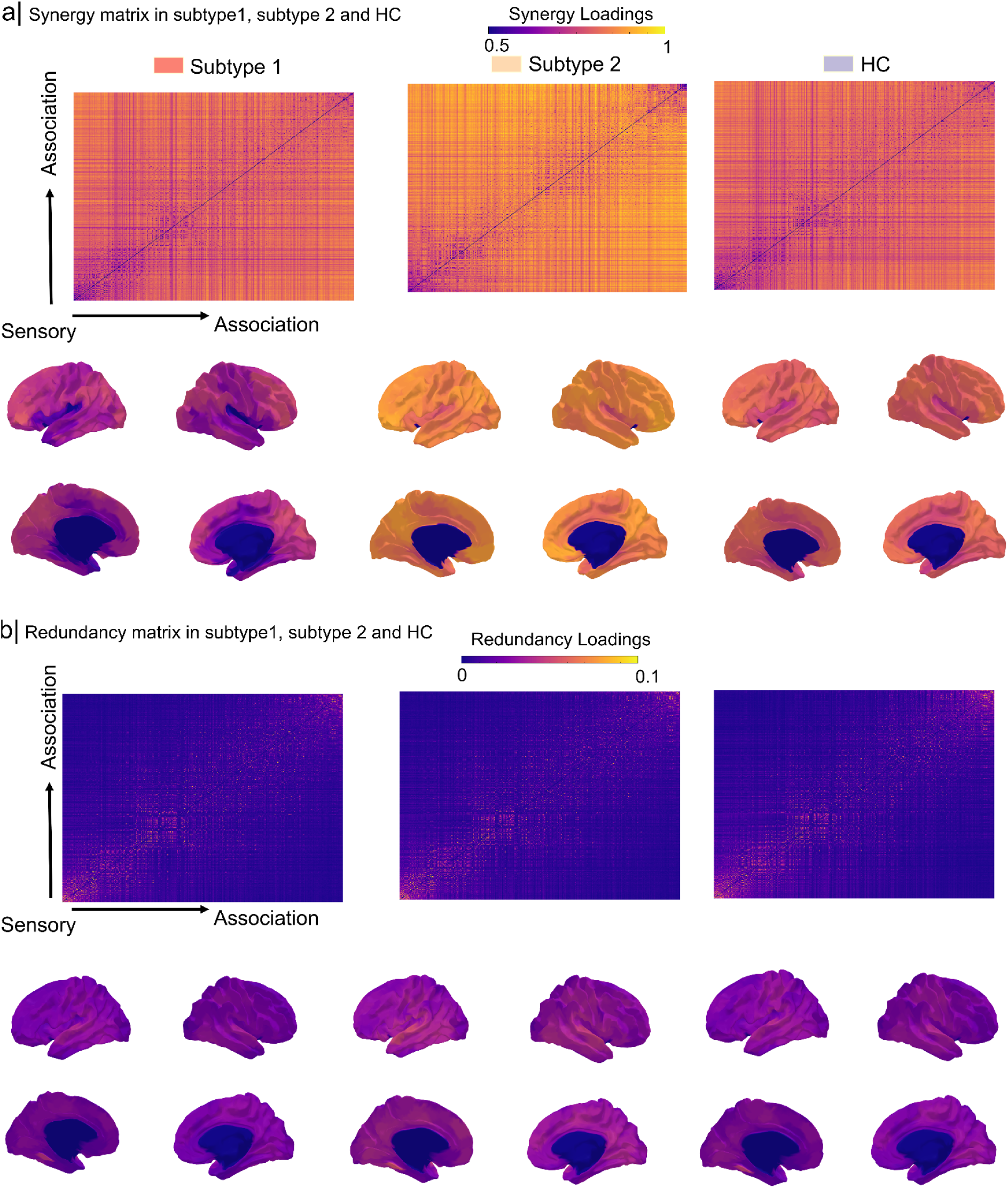
Synergy matrix and redundancy matrix subtype 1 and subtype 2. a. Synergy matrix for subtype1, subtype 2 and HC. b.Redundancy matrix for subtype1, subtype 2 and HC.

**S-figure 9.**
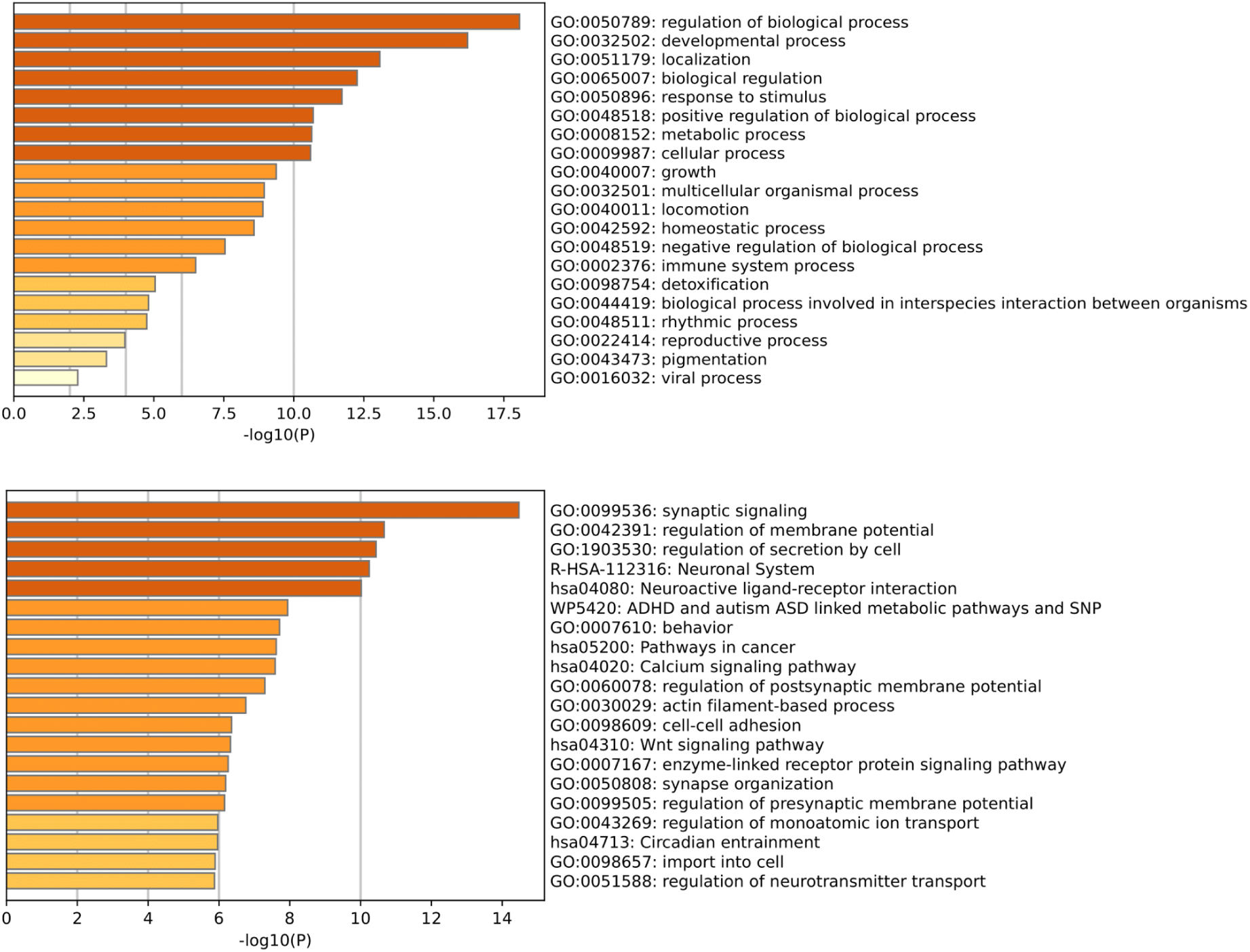
Genetic mechanism in subtype1 and subtype2 (*p_FDR_ < 0.05*).

**S-figure 10.**
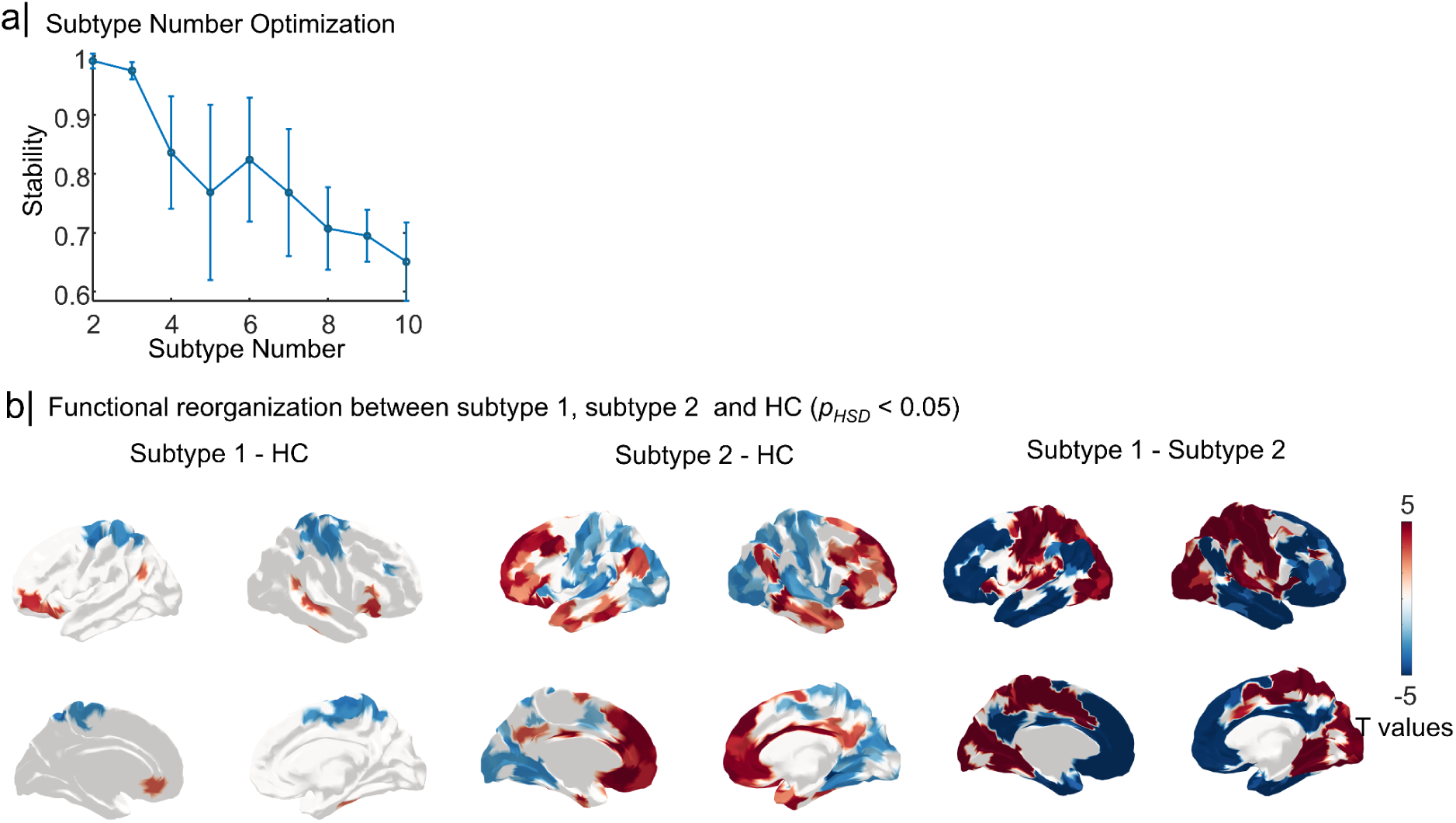
Replication results in an independent dataset. a) The optimization number is 2. b) The statistical comparison among subtype 1, subtype 2 and HC.

**S-Table 1.** Demography (HAMD means Hamilton Depression Rating Scale.

|  | Group | Number | Age | Sex (F/M) | HAMD |
| --- | --- | --- | --- | --- | --- |
| Exploration dataset | A-MDD | 258 | 15.18±1.61 | 114/144 | 24.67±5.74 |
|  | HC | 203 | 15.53±1.98 | 82/121 | ---- |
| Replication dataset | A-MDD | 73 | 16.80±1.22 | 31/42 | ---- |
|  | HC | 28 | 16.82±1.40 | 12/16 | ---- |

**S-Table 2.** Childhood Trauma Questionnaire. (CTQ, CTQ-EA means Emotional Abuse, CTQ-EI means Emotional Neglect, CTQ-PA means Physical Abuse, CTQ-PI means Physical Neglect, CTQ-SA means Sexual Abuse, CTQ-T means Total Score), HAMA means Hamilton Anxiety Scale, F means female, M means male)

|  | CTQ-EA | CTQ-EI | CTQ-PA | CTQ-PI | CTQ-SA | CTQ-T | HAMA |
| --- | --- | --- | --- | --- | --- | --- | --- |
| Score | 12.75±4.8<br>8 | 15.88±5.5<br>6 | 7.5±3.50 | 10.40±4.0<br>7 | 6.08±2.50 | 52.65±14.<br>92 | 23.65±6.3<br>2 |
| Number | 52 | 52 | 52 | 52 | 52 | 52 | 159 |

